# CellFlow enables generative single-cell phenotype modeling with flow matching

**DOI:** 10.1101/2025.04.11.648220

**Authors:** Dominik Klein, Jonas Simon Fleck, Daniil Bobrovskiy, Lea Zimmermann, Sören Becker, Alessandro Palma, Leander Dony, Alejandro Tejada-Lapuerta, Guillaume Huguet, Hsiu-Chuan Lin, Nadezhda Azbukina, Fátima Sanchís-Calleja, Theo Uscidda, Artur Szalata, Manuel Gander, Aviv Regev, Barbara Treutlein, J. Gray Camp, Fabian J. Theis

## Abstract

High-content phenotypic screens provide a powerful strategy for studying biological systems, but the scale of possible perturbations and cell states makes exhaustive experiments unfeasible. Computational models that are trained on existing data and extrapolate to correctly predict outcomes in unseen contexts have the potential to accelerate biological discovery. Here, we present CellFlow, a flexible framework based on flow matching that can model single cell phenotypes induced by complex perturbations. We apply CellFlow to various phenotypic screens, accurately predicting expression responses to a wide range of perturbations, including cytokine stimulation, drug treatments and gene knockouts. CellFlow successfully modeled developmental perturbations at the whole-embryo scale and guided cell fate and organoid engineering by predicting heterogeneous cell populations arising from combinatorial morphogen treatments and by performing a virtual organoid protocol screen. Taken together, CellFlow has the potential to accelerate discovery from phenotypic screens by learning from existing data and generating phenotypes induced by unseen conditions.

## Introduction

High-throughput perturbation screens with single-cell readouts have transformed our ability to investigate biological systems ^1–3^. Increasingly, phenotypic screens are employed to study various biological phenomena, including drug screens to identify effective therapeutic compounds ^4–6^, cytokine treatments to evaluate immune responses ^7,8^, genetic modifications to elucidate gene function ^7,9–11^, and morphogen or transcription factor manipulations for cell fate engineering ^12,13^. Such experiments can scale to hundreds or even thousands of conditions, generating large-scale perturbation atlases with rich profiles of each of millions of cells across diverse interventions ^5,8^. Beyond studying individual perturbations of one class (e.g., drug, genetics) and their combinations, it is important to understand interactions between different classes of interventions. For example, disease modeling may require examining how genetic mutations interact with drug treatments, or organoid engineering may involve multi-step protocols with sequential exposure to different media compositions and morphogen pathway modulators. Collectively, these screening approaches aim to map the relationship between diverse perturbations and phenotypic outcomes (Figure 1A).

**Figure 1.**
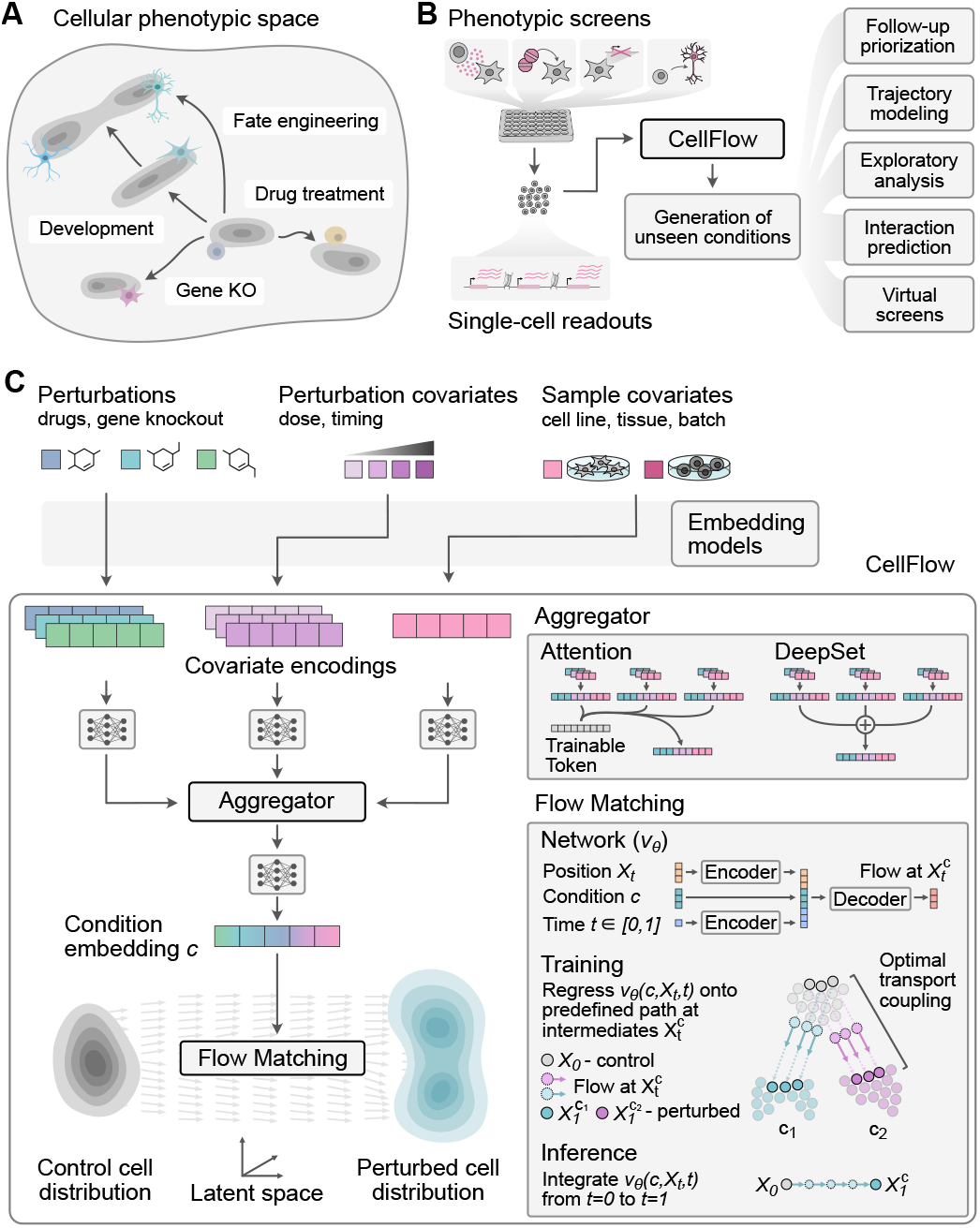
CellFlow: a tool to explore cellular phenotypic space. **(A)** Cells can change their phenotype in response to various stimuli, such as drug treatment, gene knockout or developmental cues. **(B)** Phenotypic screens help to explore the relationships between perturbation and phenotype. CellFlow can learn from such screens and generate phenotypes of unseen conditions to facilitate various downstream tasks. **(C)** CellFlow takes into account various variables of a generic experimental setup and uses aggregations strategies such as attention mechanisms and DeepSets to obtain an aggregated condition embedding. We use flow matching, which regresses predicted flows onto a predefined probability path, to learn a conditional vector field transforming a source distribution onto the perturbed distribution.

However, the complexity of biological systems and combinatorial space of potential perturbations makes comprehensive experimental characterization infeasible ^14^. Computational models can help systematically explore the phenotypic space by learning from existing perturbation data and extrapolating to uncharacterized experimental conditions. Current perturbation prediction approaches for single-cell genomics have mostly focused on predicting effects of a limited range of interventions in a limited set of differentiated cell types in 2D culture systems or isolated primary cells in suspension ^6,15–19^. However, multicellular tissues consist of heterogeneous cell populations and phenotypic screens often require sophisticated experimental setups. Comprehensive prediction of single-cell phenotypes therefore requires models that can capture heterogeneous cell distributions and incorporate information about diverse types of combinatorial interventions, such as perturbed genes, administered drugs and their dosage and timing as well as descriptors of cellular state.

To address these challenges, we introduce CellFlow, a flexible framework for modeling single-cell phenotypes induced by diverse internal or external cues. CellFlow leverages flow matching, a generative modeling technique ^20^ that has been used to generate hiqhquality samples in computer vision ^20,21^, video ^22^, and molecular design ^23,24^, in part through modeling complex distributions. We use optimal transport ^25–29^ to pair unperturbed and perturbed cells, enabling the distinction between inherent cellular heterogeneity and perturbation-induced distributional changes. To represent arbitrary types of perturbations and enable predictions for conditions out-of-distribution, CellFlow incorporates powerful pre-trained embeddings of biological entities ^30^. To model arbitrary numbers of perturbations in a permutation-invariant manner, we employ set aggregation strategies including multihead attention, a key factor to foster the success of large language models ^31^.

We demonstrate the capabilities and flexibility of CellFlow across a wide range of phenotypic screening applications. We show that it accurately models donor-specific cellular responses to cytokine treatment on a large perturbation dataset of almost ten million Peripheral Blood Mononuclear Cells (PBMCs) ^8^. To demonstrate CellFlow’s ability to model highly complex cellular distributions, we predict single-cell expression profiles across entire zebrafish embryos perturbed with different gene knockouts at various developmental stages. We show state-of-the-art performance on established perturbation prediction tasks, including prediction of gene knockout and drug treatment effects. Finally, we show CellFlow’s ability to predict heterogeneous cell populations resulting from neuron fate engineering and during organoid development. As a proof-of-principle, we perform a virtual organoid protocol screen, which identifies previously untested treatment regimens with strong effects on organoid development. Together, our results show that CellFlow can accelerate discovery from phenotypic screens by extrapolating to unseen conditions, thereby informing the prioritization of follow up experiments and enabling efficient experimental design approaches (Figure 1B).

## Results

### CellFlow is a general framework to model perturbed single-cell phenotypes

CellFlow aims to predict singlecell phenotypes under diverse perturbations by conditionally mapping a source distribution (e.g. control cells) to a perturbed population of cells. The model first encodes experimental variables and aggregates combinatorial treatments into a common condition embedding, which is then injected into the flow matching module to guide the flow from source to perturbed distributions (Figure 1C, Methods).

To accommodate a wide range of phenotypic screening scenarios, we define a generic experimental setup with three types of variables: perturbations, perturbation covariates, and sample covariates. Perturbations represent observed experimental interventions, such as drug treatments or genetic modifications. To allow predictions for interventions that were unseen during training, we employ encodings, such as molecular fingerprints ^32^ for small molecules and ESM2^30^ embeddings for the protein products of gene knockouts and proteins. Perturbation covariates represent additional modulators of treatments such as dosage or timing whereas sample covariates represent cell descriptors, independent of their treatment, such as the cell line or tissue. We refer to a combination of perturbations, perturbation covariates, and sample covariate as a single condition.

To model multiple interventions at the same time, it is necessary to encode combinations of treatments in a permutation-invariant manner. Therefore, CellFlow implements two set aggregation schemes, multi-head attention and deep sets, followed by an encoder consisting of a feed-forward network. This allows CellFlow to represent an arbitrary combination of perturbation variables in a single representation.

The condition embedding vector then serves as input into the flow matching module, which translates a source distribution to the perturbed cell population.

To learn a meaningful mapping on the population and single-cell level, we assume cells follow the least-action principle in response to a perturbation, and thus leverage optimal transport to pair samples batch-wise. We train our models end-to-end, thus learning a space of condition representations, which guide the generation of perturbed cell states from the source population. Training of flows is facilitated when the signal-to-noise ratio is high. As single-cell data can have sparse feature detection and technical variation, we leverage lower-dimensional cellular representations obtained from principal component analysis (PCA) or variational autoencoders (VAEs) ^33^.

### Scalable prediction of cytokine responses in a large phenotypic screen

To assess the predictive capabilities and scaling properties of CellFlow on high-throughput phenotypic screens, we applied it to a recent cytokine perturbation screen on almost ten million Peripheral Blood Mononuclear Cells (PBMCs) ^8^. This dataset comprises scRNA-seq samples obtained from twelve donors, each treated with each of 90 cytokines (Figure 2A). Due to their central role in the immune system, cytokines are a promising therapeutic target for numerous diseases like cancer and autoimmune disorders ^34^.

**Figure 2.**
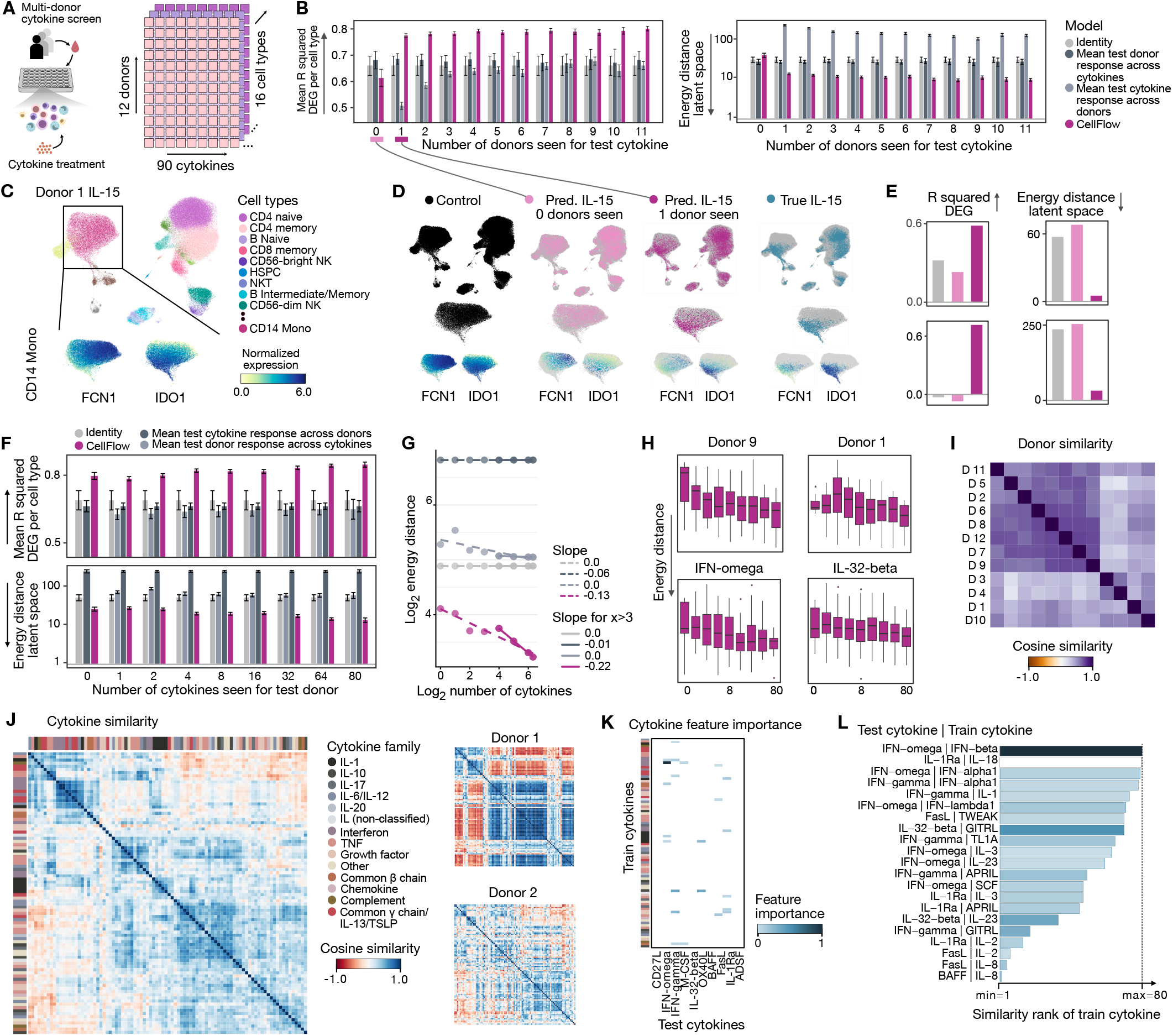
CellFlow exhibits interpretable and scalable training behaviour on 10 million PBMCs. **(A)** Peripheral Blood Mononuclear Cells (PBMCs) from twelve different donors are treated with 90 different cytokines, resulting in a dense experimental design capturing almost 10m cells. **(B)** Performance metrics for predicting donor-specific cytokine responses for varying numbers of donors for which the cytokine has been seen. The R squared of the top 50 differentially expressed genes (DEGs) per cell type is computed based on label transfer from real measurements to generated cells. One reported data point represents the mean of the R squared across cell types for a single condition (Methods). Plotted are the mean and the standard error across different training data sets with the same number of donors seen for the test cytokine (left). Analogously, the energy distance in latent space is reported for different numbers of conditions in the training data set (right, Methods). **(C)** UMAP of IL-15 treated PBMCs of donor 1, together with an inset of CD14 Mono cells colored by normalized gene expression of the top upregulated gene FCN1 and top down-regulated gene IDO1 (left). **(D)** Joint UMAP of control cells of donor 1, true IL-15 treated cells of donor 1, as well as generated cells of donor 1 treated with IL-15 from two different models. The first model was trained without IL-15 treated cells in the training dataset, while the other model included IL-15 treated cells of donor 8 in the training data set. Each column shows one of the four cell populations highlighted (top), together with an inset of CD14 Mono cells (middle), and the normalized gene expression of FCN1 and IDO1 (bottom). **(E)** Quantification of the visual results of the identity model (i.e. using the control distribution of donor 1 as predicted IL-15 treated cells), CellFlow trained with no IL-15 cells seen, and CellFlow trained with IL-15 treated cells from donor 8. **(F)** Performance of CellFlow and baselines for predicting the cytokine response of a new donor based on a varying number of cytokines measurements for the new donor. Mean and standard error are computed analogously to (b). **(G)** Linear fits of the mean energy distances displayed in (g) in log-log space. Slopes are computed from models trained on at least one cytokine for the test donor (dashed line) and trained on at least 16 cytokines for the test donor (solid line). **(H)** CellFlow’s performance metrics filtered for predicted populations specific to donor 9 (top left) and donor 1 (top right), as well as specific to IFN-omega (bottom left) or IL-32 beta (bottom right). **(I)** Donor similarities computed from responses across all cytokines (Methods). **(J)** Cytokine similarities computed from responses across all donors (left), and from responses of donor 1 (top right) and donor 2 (bottom right, Methods). **(K)** Importance of presence of cytokine in training data set for a good performance of CellFlow based on the coefficients of a regularized linear model (Methods). **(L)** Ordered enumeration of non-zero importances scores displayed in (K).

We first investigated CellFlow’s predictive performance in relation to the number of donors for which a cytokine treatment has been observed. For a given test combination of a donor and a cytokine, we systematically varied the number of donors treated with that cytokine included in the training data. For each number of donor-specific cytokine treatments included in the training data, we held out different sets of conditions during training (Methods). We evaluated the generative performance using the energy distance between the true and the predicted cell population in PCA space (Methods). To evaluate predicted gene expression, we transferred cell type labels of generated cells from true ones using the one-nearest neighbor classifier, and computed the R squared between the means of the true and the predicted gene expression for the top 50 cell type- and condition-specific differentially expressed genes (DEGs, Methods). To assess the ability to learn donor-specific cytokine responses beyond simple assumptions, we compared the performance against three baseline models: (1) an identity model assuming no effect (identity), (2) a model assuming a cytokine treatment has consistent effects across donors (donor mean), and (3) a model assuming all cytokine treatments have the same effect within a donor (cytokine mean). We found that when no samples of a test cytokine treatment were included in the training set, CellFlow was not able to predict a cytokine’s donor-specific response more accurately than baseline methods (Figure 2B, Figure S1A). However, as soon as the cytokine response of one donor was included in the training data, the performance drastically improved (mean DEG R-squared of 0.76 vs. 0.60), outperforming all baselines (Figure 2B). This performance increase can also be illustrated in a UMAP embedding ^35^ (Figure 2C, Methods). While the donor-specific cell population generated without having seen IL-15 for any other donor resembled the control cells, having seen IL-15 for only one other donor shifts predictions towards the true distribution, both on a population level and in terms of generated expression of the differentially expressed genes (Figure 2D,E). These results suggest that the ESM2 representation of the cytokine is not sufficient for CellFlow to extract the functional effect on gene expression level but it was able to calibrate to donor-specific effects as soon as the cytokine treatment effect has been measured for one donor. Moreover, measuring the cytokine treatment for multiple donors still increased the performance of CellFlow, but with decreasing marginal effects. Thus, given a sparse budget to explore the donor-specific responses of the remaining cytokine treatments, the most promising way to obtain reliable predictions is prioritizing the number of different cytokine treatments measured rather than the number of donors for a single treatment.

Next, we assessed the prediction of cytokine effects for new donors. To obtain a representation of each donor, we computed the donor-specific mean gene expression vector of the control population. This naive representation was sufficient for CellFlow to predict cytokine-specific donor responses more accurately than any baseline model (Figure 2F). In practice, it can be feasible to measure a limited number of cytokine treatments for a new donor. Therefore, we evaluated the performance of all models across different numbers of cytokine treatments measured for a new donor, which showed that adding more cytokine treatments further increased performance resulting in an almost 10-fold improvement over baselines (Figure 2G and Figure S3A). We further tested whether CellFlow could learn donor-specific responses. Analysis revealed positive correlations between predicted and true donor similarities based on test cytokine responses, although CellFlow tended to underestimate the magnitude of these differences between donors (Figure S2C-E).

We were able to observe a clear scaling relationship between CellFlow’s performance and the number of seen conditions. This scaling law followed a linear fit in loglog space with an intercept of −0.22 and an R squared value of 0.999 for 16, 32, 64, and 80 cytokines included in the training data (Figure 2G, Methods). Scaling behavior differed across individual donors and cytokines (Figure 2H). For example, CellFlow’s performance rapidly increased for donor 9. In contrast, the performance for donor 1 only increased when including a large number of treatments (Figure 2H). This may be explained by the high similarity of donor 9 to other donors, while donor 1 was highly dissimilar to all other donors (Figure 2I and Figure S2A). Analogously, we found a similar distinction between the scaling curves for IFN-omega and IL-32-beta (Figure 2I), suggesting the presence of a similar cytokine as IFN-omega and the lack of a cytokine similar to IL-32-beta. Indeed, IFN-omega had a highly similar effect as IFN-beta (cosine similarity=0.97), while the most similar cytokine to IL-beta only had a cosine similarity of 0.55 (Figure 2J, Figure S2B, Methods). The dissimilarity led us to ask whether CellFlow was able to generate such a novel cell state. Indeed, we found IL-32-beta generations to be closer to the true set of cells than any population in the training set for 9 out of 12 donors (Figure S3B). An analogous analysis revealed that predicted populations are highly donor-specific (Figure S4B). To further investigate this scaling behavior, we assessed the relationship between predictive performance and the presence of certain cytokines in the training dataset. For this, we computed feature importance scores for each test cytokine with respect to all other cytokines using a regularized linear model (Methods). We observed that prediction accuracy for IFN-omega strongly depends on the presence of IFN-beta in the training data, which is consistent with the similar cellular responses elicited by these two cytokines. Similarly, the majority of identified train cytokines which positively influence CellFlow’s performance could be explained by a high similarity to the test cytokine (Figure 2K). Together, this shows that CellFlow exhibits scaling behaviour with respect to the number of measured cytokines, which can be explained by similarity relationships between cytokines and donors.

In summary, having trained 792 CellFlow models on up to ten million cells, resulting in more than 50 million generated cells (Methods), we found that CellFlow is stable to train and capable of accurate predictions exceeding baselines in almost all settings. We also observed a consistent and explainable training behaviour, satisfying scaling laws in the number of seen conditions.

### Modeling the perturbed development of zebrafish embryos

To understand CellFlow’s ability to model perturbations of developing multicellular systems, we investigated the predictive performance on a phenotypic screen from an entire developing organism. We used ZSCAPE ^10^ (zebrafish single-cell atlas of perturbed embryos), which measured the effects of 23 gene knockouts induced through F0 genome editing measured on scRNA-seq profiles across five points during development (Figure 3A). Here, each knockout was only profiled at a subset of the five considered time points. Thus, we tested whether CellFlow could accurately predict the cellular phenotypes of perturbed developing embryos at unobserved developmental stages. We simulated this setting by predicting the cellular states of mutants, resulting in 71 trained models, each corresponding to one held-out condition defined by a combination of knockout and time point. We considered three baseline models: (1) An identity model assuming a gene knockout has no effect (identity); (2) the mean effect of all other perturbations observed at the same time point; and (3) the mean effect of the same gene knockout at the remaining observed time points (Methods). CellFlow outperformed all baselines across different metrics (2x improvement in energy distance; Figure 3B, Figure S4A). To assess predictions on the cell type level, we transferred cell type annotations from measured cells to generated cells (Methods). We found that the performance was consistently above baseline across gene knockouts, developmental stage of the embryo, perturbation strength as well as time point interpolation and extrapolation scenarios (Figure 3C-F, Figure S4B). However, the extent of improvement decreased with more mature developmental stages of the embryo, possibly due to fewer phenotypes being observed at later time points and the increasing heterogeneity of the embryos. These evaluations show that CellFlow is able to learn meaningful gene knockout-specific developmental phenotypes.

**Figure 3.**
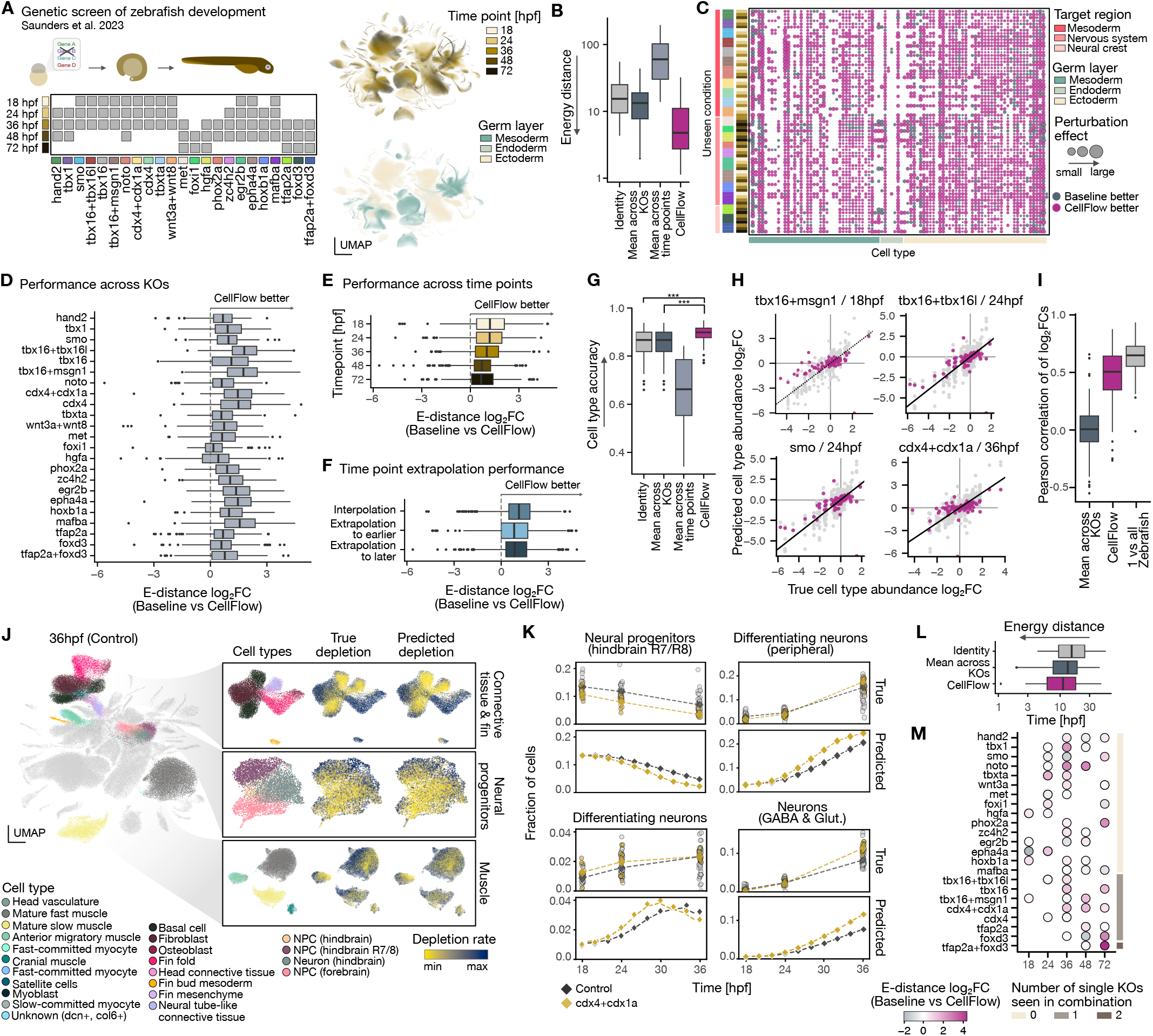
Organism-scale prediction of perturbed zebrafish development. **(A)** ZSCAPE7 (zebrafish single-cell atlas of perturbed embryos) captures the developing zebrafish at developmental stages ranging from 18 hpf (hours post fertilization) to 72 hpf with different genes knocked out. **(B)** Energy distance in latent space for CellFlow, as well as three baseline models for the task of predicting the cellular phenotype of a perturbed zebrafish at a certain time point: The identity assumes no perturbation effect, while the mean models assume either a constant effect across gene knockouts or a constant effect across time points. **(C)** Dotplot showing CellFlow’s performance relative with the knockout mean baseline per cell type and time point. Cell types are categorized according to germ layer (bottom strip), and ordered by similarity (Methods), while gene knockouts are ordered by target region, gene knockout and developmental time (left strips). The size of the dot indicates the effect of the perturbation on the specific cell type, measured with the energy distance. **(D)** Comparison of CellFlow’s performance (measured by energy distance per cell type) with the best baseline model aggregated across cell type and developmental time. **(E)** Performance gain by CellFlow with respect to the best baseline model aggregated across cell type and gene knockout. **(F)** Performance gain of CellFlow with respect to the best baseline model aggregated to the relative temporal position of the left perturbed developmental stage, categorized into interpolation (there are measurements for the same genetic knockout in both and earlier and a later time point), extrapolation to earlier time point (there is no measurement of the same genetic knockout in an earlier developmental stage), and extrapolation to a later time point (no measurement of the same genetic knockout in a later developmental stage). **(G)** Accuracy of predicted cell type proportions after perturbation of CellFlow and the three baseline models. ^*^<10-2, ^**^<10-3, ^***^<10-4, unpaired t-test. **(H)** True and predicted log fold changes of cell types for a certain perturbation and developmental time point of the zebrafish. Gray dots correspond to estimates of single perturbed zebrafish (as opposed to the union of cells across zebrafish of same perturbation and developmental stage, which is taken as the ground truth), providing a notion of noisiness of the data. **(I)** Pearson correlation between true and predicted log-fold changes across all 71 predicted perturbed states of the developing zebrafish, together with a reference obtained from log-fold changes of a single perturbed zebrafish with respect to the union of all perturbed zebrafish. **(J)** UMAP of control cells of a zebrafish at developmental stage 36hpf, with all cell types colored which depletion rates are calculated for (left, Methods). Insets visualize true and predicted depletion rates, which are computed from true cdx4/cdx1a-perturbed cells and predicted cdx4/cdx1a-perturbed cells at time point 36hpf. **(K)** Cell type proportions of selected cell types of the Central Nervous System at different developmental stages. The top boxes show the cell type fractions of single control zebrafish (gray dots) and single perturbed zebrafish (yellow dots), as well as the fraction of the union of all control zebrafish (dark gray diamond) and all perturbed zebrafish (yellow diamond). The dotted line depicts a linear interpolation between the time points. The bottom boxes show predicted cell type proportions for control and perturbed zebrafish, interpolated to densely sampled time points which have not been measured in the dataset. The model has been trained with the cdx4/cdx1a perturbed population only present at 18hpf and 36hpf, i.e. 24hpf was not included in the training data. **(L)** Performance of CellFlow and baseline models for the task of predicting perturbed zebrafish without having seen the same genetic perturbation at any time point. **(M)** Improvement of CellFlow with respect to the best baseline model, which models the perturbation as the mean displacement of all other perturbations at the same time point.

One major phenotypic impact of the knockouts is relative changes in cell type proportions in the organism. Thus, we quantified CellFlow’s ability to accurately predict changes in cell type proportions. We found that CellFlow accurately predicted cell type proportions based on transferred annotations (mean accuracy=0.89 vs. 0.85 of best baseline; Methods), outperforming all baselines (Figure 3G). To understand to what degree changes in cell type abundances were predicted, we compared true and predicted fold changes. We found predictions were generally correlated with true proportion changes (mean correlation=0.43) and accurately captured whether cell type abundance was increased, decreased or not affected (Figure 3H,I). However, strong effects were often underestimated (Figure 3H). This overly conservative behavior may in part be due to the variability of individual mutants (Figure 3H,I) and the high variability of the one-nearest neighbor classifier used for cell type annotation transfer (Figure S4B). To further assess CellFlow’s predictions on the single-cell level, we computed depletion rates of single cells induced by the combined knockout of cdx4 and cdx1a, as previously investigated ^10^ (Methods). We used our model to extrapolate the effect of the genetic perturbation observed at 18hpf and 24hpf to time point 36hpf and found cell-specific true and predicted depletion rates to be highly correlated across affected organs of the zebrafish, even if the magnitude was often underestimated (Figure 3J, Figure S4C-E). For example, given observations in earlier time points, CellFlow was able to project the impact of the perturbation in hindbrain neural progenitor cells to 36h. This analysis demonstrates that CellFlow can accurately capture both organismwide and tissue-specific cell type abundance changes, enabling the prediction of perturbed cell proportions for unobserved time points.

We next leveraged CellFlow to model the continuous, perturbed development of the central nervous system (Methods). We held out the cdx4/cdx1a mutant at time point 24, allowing us to compare the predicted cell type composition of the central nervous system with the true, unseen one. CellFlow’s predictions of cell type fractions accurately matched the mean of the ground truth distribution for most cell types despite the high variance between individual zebrafish (Figure 3K, Figure S4F). This allowed us to further generate cells corresponding to a control or cdx4/cdx1a perturbed zebrafish for densely sampled time points, resulting in an interpolation of cell type abundance changes during development.

A more challenging task than predicting the effect of gene knockouts which have been seen at other developmental stages is modeling the evolution of an embryo with a completely unseen genetic perturbation. This requires CellFlow to fully rely on the discriminative power of the gene knockout embedding (here ESM2). While CellFlow performed above baseline on average, it did so less consistently than in the previous task (Figure 3L,M, Figure S4G). Analogously to the prediction of effects of new cytokines on PBMCs (Figure 2B), this suggests that the low number of observed gene knockouts in the training data makes it difficult to learn distinctive patterns in the data.

### Predicting perturbation effects for diverse and complex experimental designs

We next asked how CellFlow would perform on established perturbation prediction benchmarks across diverse experimental designs. We first evaluated its ability to predict single-cell expression responses to drug treatments in cancer cell lines ^4^ (sciPlex3 dataset; Figure 4A), comparing against established perturbation prediction methods chemCPA ^16^, biolord ^19^, and CondOT ^17^ (Figure 4B, Figure S6A). CellFlow performed competitively across all metrics, with particularly strong performance in predicting high-dose drug effects. For lower dosages, which typically induce more subtle expression changes, all methods showed comparable performance, with none significantly outperforming a simple identity baseline (Figure S6A). To illustrate CellFlow’s ability to model continuous dose-response relationships, we examined predictions for MCF7 cells treated with Dacinostat, a drug which had not been seen during training. The model correctly captured monotonic changes of differentially expressed genes in response to increasing drug concentration while maintaining realistic cell states close to the phenotypic manifold (Figure 4C, Figure S5A,B). Moreover, CellFlow’s inherent optimal transport modeling approach allowed to trace the response of single cells across different dosages, overcoming the limitation of the destructive nature of sequencing technologies (Figure 4C, Figure S5C,D). We further leveraged the sciPlex3 dataset to compare PCA and VAEs as possible methods for encoding and reconstruction of cells. We computed upper bounds of CellFlow’s performance by encoding and subsequently decoding held-out conditions independently of CellFlow. While we found PCA encoders to be more powerful with respect to distributional metrics, the VAE was more powerful at fitting the mean gene expression (Figure S5E, Methods).

**Figure 4.**
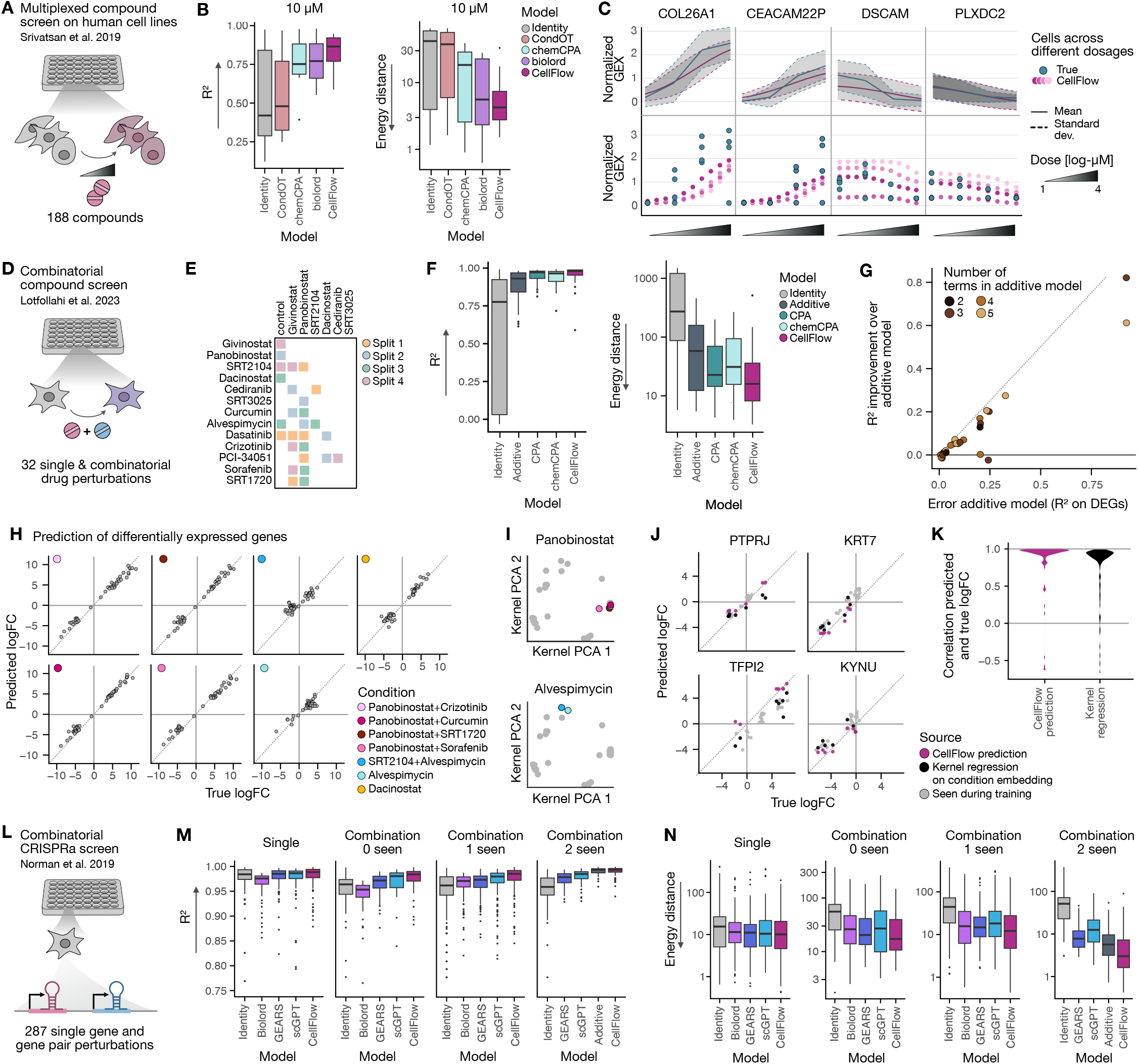
CellFlow outperforms other methods on diverse perturbation prediction tasks. **(A)** The sciPlex3 dataset contains measurements of three cancer cell lines treated with various drugs at different dosages. **(B)** Performance of CellFlow and other established methods for the task of predicting the effect of unseen drugs of the highest dosage on all three cell lines, measured by R squared in normalized gene expression space as well as with the energy distance in latent space (Methods). **(C)** True and predicted gene expression of differentially expressed genes of MCF7 cells treated with Dacinostat, a drug which has not been seen during training, across different dosages. CellFlow enables tracing the gene expression of single cells (colored by different shades of magenta) across different interpolated dosages (bottom). In contrast, gene expression of measured cells is only available for four different dosages and cells are unpaired across different dosages. **(D)** The combosciplex dataset captures gene expression responses of the A549 cell line treated with different drugs. **(E)** Combinations of drugs present in the combosciplex dataset colored by test split. **(F)** R-squared of normalized gene expression aggregated across all four splits for CellFlow and other established methods (Methods), as well as the energy distance in latent space (Methods). **(G)** Comparison of the performance of CellFlow and the additive model with respect to the R-squared of treatment-specific differentially expressed genes. **(H)** True and predicted log-fold changes of treatment-specific differentially expressed genes for all drug perturbations in split 3. **(I)** Learned embedding space by CellFlow with all conditions in gray which have been seen during training, and all test-conditions containing Panobinostat (top) or Alvespimycin (bottom) colored. **(J)** Predicted log-fold changes induced by treatments for lung cancer suppressing genes PTPRJ, KRT7, TFPI2, and KYNU. Predictions are either obtained from generated gene expression CellFlow (magenta) or from a linear model trained on CellFlow’s learnt embedding space (dark gray). **(K)** Correlations between true and predicted log-fold changes across all conditions and all four cancer suppressors PTPRJ, KRT7, TFPI2, and KYNU for predictions obtained from generated samples, and predictions obtained from a linear model on CellFlow’s learnt embedding space. **(M)** The genetic perturbation dataset captures cellular responses to genetic perturbations yielding overexpression of genes on the K562 cancer cell line6. **(N)** Comparison of performance of different methods based on R squared of normalized gene expression and energy distance in latent space (Methods).

We next assessed CellFlow’s ability to predict the effect of drug combinations using the combosciplex dataset, which contains scRNA-seq data for 31 single or combinatorial drug treatments on the A549 lung cancer cell line ^6^. We split the data such that each of the individual drugs in the test data has been observed in a different combination in the training data (Figure 4E). CellFlow outperformed other established methods as well as an expression-space additive model (Figure 4F-G, Figure S6B, Methods). As recent studies have highlighted the difficulty of beating simple additive baselines ^6,36^, we focused on the comparison between CellFlow and the additive baseline. As not all drugs appearing in combinations were measured individually, the additive model sometimes required more than two terms (Methods). We found that CellFlow largely outperformed the additive model, independent of the number of additive terms, and reduced the error of the additive model by 66% on average as measured by R squared values for DEGs (Figure 3G). CellFlow effectively recovered log-fold changes of perturbation-specific DEGs, showing particular strength in predicting large-magnitude changes while struggling more with subtle expression shifts (Figure 4H), which is consistent with our observations in the sciPlex3 dataset.

Following recent interests in learning representation of drugs with foundation models ^37^, we analyzed whether CellFlow learnt a meaningful functional embedding of the combinations of drugs. Visually, we found similar combinations of drugs to be close in embedding space (Figure 4I). We quantified the biological meaningfulness of the learnt latent space by predicting the log-fold change of gene expression from the learnt embedding. In particular, we modelled the response of TFPI2 and PTPRJ, whose overexpression is associated with better clinical outcome in lung cancer patients ^38,39^, and KRT7 and KYNU whose overexpression results in poor prognosis ^40,41^. We found that a linear predictor trained on the learnt embedding space produced meaningful predictions, but did not match the accuracy of predictions directly computed from generated cell profiles (Figure 4J,K).

To investigate CellFlow’s ability to predict the effect of genetic perturbations, we evaluated its performance on a dataset capturing the response to gene overexpression ^9^ (Figure 4L). Following a previously established evaluation holdout strategy for this dataset ^18^, we assessed predictions on three categories of held-out conditions: previously unseen single genes, combinations where all genes were unseen during training, combinations containing one unseen gene, and novel combinations of genes that had been individually observed. We benchmarked CellFlow against GEARS, biolord, and scGPT ^42^, as well as the identity baseline. CellFlow achieved the highest median R-squared and the lowest median energy distance across all evaluation splits (Figure 4M,N, Figure S6C, Methods). In line with drug perturbation modeling as outlined above, we defined an additive model for the group of conditions comprising unseen combinations of seen genes. This additive model performed comparably to CellFlow when evaluating mean gene expression (0.99 mean R-squared for both models), but in terms of energy distance, CellFlow improved upon the additive model by 2-fold (Figure S6C).

To further showcase CellFlow’s ability to address complex experimental designs, we used a recent study of genetic perturbations on six different cancer cell lines stimulated with five different cytokines ^7^. In this dataset, all cell lines were treated with each of the cytokines, but the each cytokine treatment was combined with a different set of gene knockdowns (Figure S7A). As with the PBMC and zebrafish examples, other methods were insufficient to capture this experimental design. We therefore compared CellFlow against two baselines: an identity model assuming no perturbation effect, and a model predicting the mean effect of all other perturbations in the same cell line/pathway combination (Methods). CellFlow is competitive with the additive model with respect to the performance on DEGs across perturbation strengths (Figure S7B, Methods). We then tested whether CellFlow could predict perturbation effects across cell lines (Figure S7C, Methods). Again, CellFlow clearly outperformed baseline models for strong effects, while for subtle expression changes, the identity model often yielded better predictions (Figure S7D). Based on previously described conserved and context-specific perturbation programs ^7^, we evaluated CellFlow’s predicted effect on curated sets of target genes on the interferon-gamma (IFNG) treated BXPC3 cell line, a condition manifesting strong and heterogeneous perturbation effects ^7^. CellFlow’s predictions were more similar to the ground truth than the responses of any other cell line or the mean thereof (Figure S7E). As expected, effects of cell line-specific perturbation responses were harder to predict than those that are conserved across cell lines (Figure S7E). Finally, we assessed whether CellFlow could predict gene knockdowns to a completely unseen cell line, a challenging task as the performance relies on CellFlow extracting patterns from only five cell line embeddings (Figure S7F). While predictions were better than the ones obtained from baseline models for strong perturbation effects, CellFlow was not able to outperform the identity model for smaller effects (Figure S7G).

The flexibility of CellFlow does not only apply to the choice of experimental designs, but also to the type of readout. As a proof of concept, we demonstrated its capability for predicting the effect of chemical perturbations on cancer cell lines measured by the 4i technology ^17,43^ (Figure S8). Altogether, we demonstrated CellFlow’s performance on a large variety of perturbation experiments, allowing for more flexibility than previous methods which are mostly tailored towards specific experimental setups.

### Modeling neuron programming through combinatorial morphogen treatment

We next sought to use CellFlow to predict the outcome of cell fate programming experiments where conditions can generate unexpected and novel cell states. For this task, we leveraged a dataset based on a combinatorial morphogen screen on NGN2-induced human neurons (iNeurons) ^13^. Such *in vitro* systems are challenging to model, as each condition can contain heterogeneous cell fates in response to complex interactions of intrinsic and extrinsic signalling modulators. In this screen, NGN2 expression was induced in iPSCs, followed by treatment with morphogen pathway modulators during neuronal differentiation (Figure 5A). The treatment conditions comprised combinations of modulators of anterior-posterior (AP) patterning (RA, CHIR99021, XAV-939, FGF8) with modulators of dorso-ventral (DV) patterning (BMP4, SHH), each applied in multiple concentrations. scRNA-seq readouts of patterned NGN2-iNs showed that treatments resulted in diverse distributions of neurons within each condition, including three distinct neuron classes and various brain region identities such as forebrain, midbrain, hindbrain and spinal cord as well as peripheral sympathetic and sensory neurons (Figure 5B,C).

**Figure 5.**
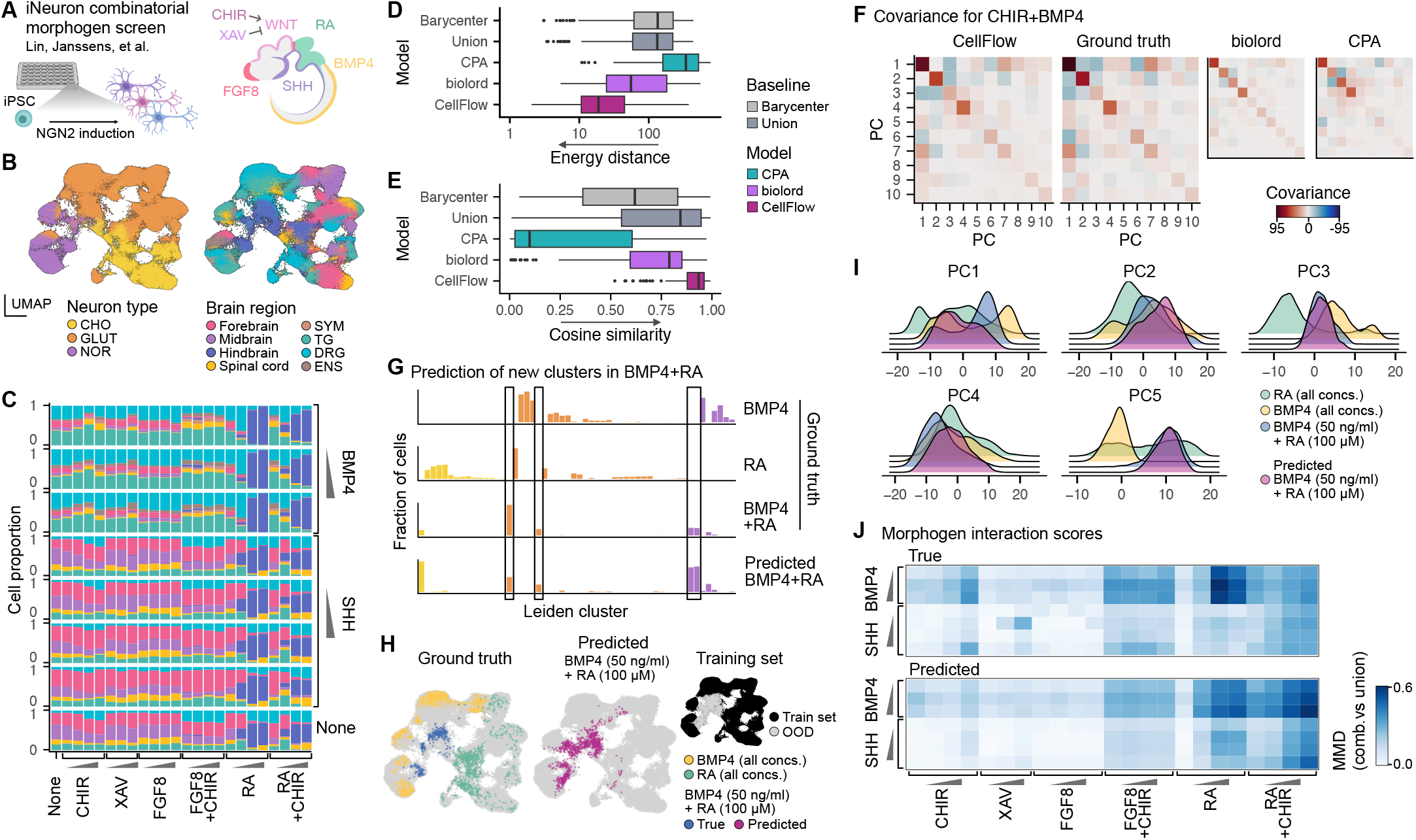
Neuron fate prediction from combinatorial morphogen treatment to enable condition prioritization. **(A)** Schematic of the experimental setup for the iNeuron combinatorial morphogen screen. iNeurons were treated with morphogen pathway modulators during neural differentiation from pluripotency. **(B)** UMAP embedding of scRNA-seq data from all screening conditions colored by neuron type (left) and brain region identity (right). **(C)** Bar plot showing the proportion of brain regions in individual screen conditions. **(D)** Boxplot showing energy distance between predicted and true cell distributions for baselines, established perturbation prediction methods and CellFlow. **(E)** Boxplot showing cosine similarity of true and predicted leiden clusters. **(F)** Heatmap showing covariance between the first ten principal components for the true CHIR+BMP4 condition, as well as CellFlow, biolord and CPA predictions. **(G)** Bar plot showing the leiden cluster proportions for the true BMP4, RA and BMP4+RA condition as well as the BMP4+RA condition predicted by CellFlow. **(H)** UMAP embedding showing the cells belonging to the true BMP4, RA and BMP4+RA conditions (left), BMP4+RA cells predicted by CellFlow projected onto the UMAP embedding (middle) and UMAP embedding showing the cells included in the training set (right). **(I)** Density plots showing marginal distributions over the first five principal components for the true BMP4, RA and BMP4+RA condition as well as the BMP4+RA condition predicted by CellFlow. **(J)** Heatmap showing the maximum mean discrepancy between cell distributions arising from combinatorial morphogen treatment and the union of distributions from individual morphogen treatments.

To assess the ability of our model to predict neuron distributions induced by combinations of pathway modulators, we evaluated prediction performance on held-out combinatorial treatments. Specifically, in each training run, we withheld all experimental conditions containing certain combinations of AP and DV modulators (Methods), then assessed the ability to predict cellular states under these unseen treatment combinations. We compared the performance of CellFlow with CPA and biolord as well as two baseline models: the union of cellular distributions of individual treatments and their distributional mean (Wasserstein barycenter ^44^). Intuitively, these baselines represent cases where combinatorial treatments generate a combination of states observed under individual treatments and an intermediate cellular state, respectively. We found that CellFlow strongly outperformed both existing methods and baselines in terms of energy distance and other distributional metrics (≥ 2.5x mean improvement in energy distance; Figure 5D and Figure S9A-C). We additionally sought to understand whether less accurate predictions default to predicting familiar training examples or generate states outside of the training distribution. While predictions were generally close to training conditions, they did not move closer as prediction error increased (Figure S9D, left). Instead, higher prediction errors correlated with greater divergence from the average training distribution (Figure S9D, right), indicating that CellFlow’s errors tend to occur when attempting to model new cellular states rather than from incorrectly defaulting to known states.

To better evaluate the accuracy of predicted cell identity composition, we compared true and predicted Leiden cluster abundances, obtained by nearest neighbor-based label transfer ^45^ (Methods). This revealed that predictions achieved consistently high cosine similarity scores (mean=0.91) with true cluster distributions, exceeding the performance of other methods and baselines (≥ 70% mean improvement; Figure 5D,E). CellFlow also accurately captured the covariance structure of cell state distributions, demonstrating its ability to model realistic cell populations (Figure 5F and Figure S9E). These results demonstrate that previous methods for perturbation prediction struggle to capture heterogeneous phenotype distributions that arise during fate programming experiments and often do not surpass simple baselines, highlighting the power of CellFlow to model cellular state transitions.

The combination of BMP4 and RA resulted in new cell states that were not observed in either individual treatment ^13^. To assess whether such interactions between pathway modulators could be captured by CellFlow we compared the distribution of true and predicted cells across Leiden clusters, which revealed that CellFlow accurately predicted emergent populations of cells (Figure 5G). We further visualized these predictions on a UMAP projection (Methods), illustrating that CellFlow can predict new cell populations that were not observed during training (Figure 5H). This is also reflected in the marginal distributions along individual principal components, which show that CellFlow correctly learns an intermediate distribution which deviates from distributional averaging or addition of individual treatment effects (Figure 5I and Figure S9F).

Given CellFlow’s ability to accurately predict combinatorial treatment outcomes, we explored its utility in identifying morphogen interactions resulting in emergent new phenotypes, thereby enabling the prioritization of experimental follow-ups. To measure the degree to which morphogen combinations induce new cell states, we devised an interaction score based on the maximum mean discrepancy between combinatorial treatment effects and the union of distributions from individual treatments (Methods). When comparing these interaction scores between predicted and experimental data, we found that CellFlow successfully captured the overall pattern of interactions, particularly identifying combinations with high interaction scores such as BMP4+RA and BMP4+FGF8+CHIR. However, we observed discrepancies in certain combinations, such as SHH+XAV and SHH+CHIR, where interaction strength was highly concentration-dependent (Figure 5J). Despite these limitations, these results indicate CellFlow’s potential as a tool for guiding experimental design by predicting which combinations of pathway modulators are likely to yield novel and unexpected cellular states.

### A virtual organoid protocol screen

*In vitro* organoid models of brain development are complex tissues that provide inroads to study the mechanisms underlying human patterning. We applied CellFlow to model the cell type composition of human brain organoids induced by multi-step organoid protocols ^46^. To cover diverse organoid protocols, we assembled three scRNA-seq datasets of human brain organoid morphogen screens ^47–49^ for training and evaluation. In these screens, organoids were grown from pluripotency and exposed to different morphogen pathway modulators at various timings during development. Across the three datasets, 23 pathway modulators were applied in various combinations at defined time windows between day 1 and day 36 of development, comprising a total of 176 conditions. To train the model jointly on all datasets, we harmonized protocol configurations into a common protocol encoding, capturing the pathway modulators, their respective concentrations, timings and information about the modulated pathway (Figure 6A,B, Methods). This representation also incorporated a one-hot encoding of the dataset label, to account for other dataset-specific experimental parameters (base protocol), such as media composition or usage of matrigel. To integrate scRNA-seq data of the three datasets and allow for comparison of organoid cell states with primary counterparts, we projected scRNA-seq data from all datasets to a single-cell transcriptomic atlas of the developing human brain ^50^ using scANVI ^45,51^ (Methods). This joint latent space allowed us to annotate brain region and cell types across datasets through nearest neighbor-based label transfer, revealing the broad distribution of brain regions generated by the diverse brain organoid culture conditions (Figure 6C and Figure S10A).

**Figure 6.**
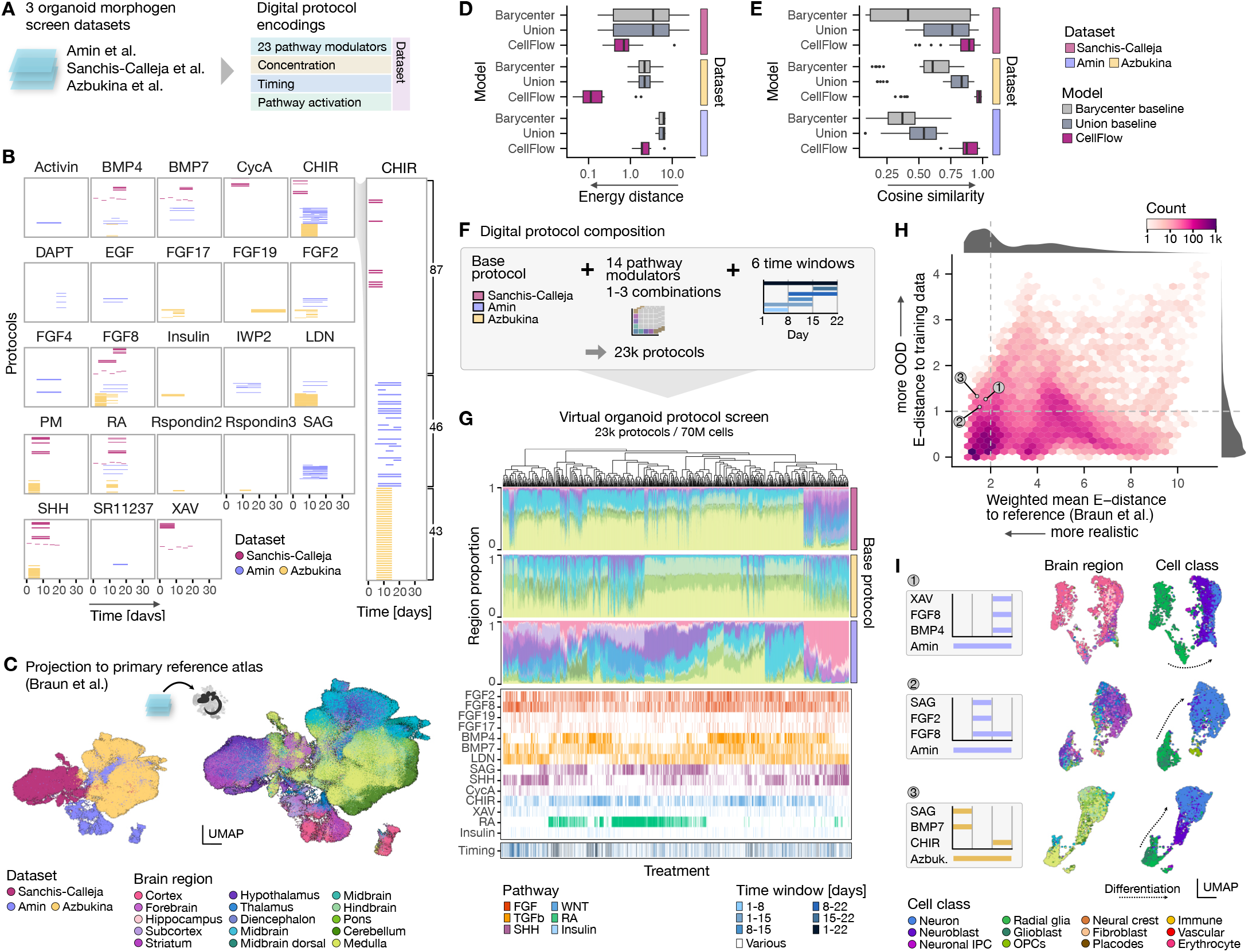
Integrated learning from systematic morphogen experiments enables virtual neural organoid protocol screening. **(A)** Schematic of integrating three organoid morphogen screen datasets to obtain consistent digital protocol encodings. **(B)** Representations of morphogen timings in all protocols across the three datasets. **(C)** UMAP embedding obtained by projecting scRNA-seq data of all three datasets onto a primary reference atlas45 using scANVI46. **(D and E)** Box plots showing the prediction performance on held out morphogen combinations in terms of energy distance of distributions **(D)** and cosine similarity of predicted leiden clusters **(E). (F)** Schematic of digital protocol composition by combining a base protocol with combinatorial pathway modulator treatments during defined time windows, resulting in 23k virtual protocols. **(G)** Predictions of the virtual protocol screen. Bar pot showing brain region composition of all predictions (top) as well as morphogen usage (middle) and timings (bottom). **(H)** Density scatter plot showing weighted mean energy distance to the reference atlas (measure of realism) versus energy distance to the closest training condition (measure of novelty) for all predictions. **(I)** UMAP embeddings showing examples for three predictions resulting in different brain region-specific progenitor and neuron populations. Colors indicate brain region (left) and cell class (right).

To assess the ability of CellFlow to learn from these systematic screens to predict cell type distributions for new protocol configurations, we first evaluated its performance on held out combinatorial treatments. We withheld all conditions containing certain combinations of pathway modulators during training, then assessed predictions for these unseen treatment combinations (Methods). Analogously to the evaluation on iNeurons, we compared the performance of CellFlow against distributional baselines (Wasserstein barycenter, union) based on energy distance and cosine similarity of predicted leiden clusters (Figure 6D,E). CellFlow achieved low energy distances and high cosine similarity scores (median=0.94), largely outperforming baselines (5.7x mean improvement in energy distance over union, Figure S10B,C). Corroborating our results from iNeurons, we found that the model was able to capture the interaction between BMP7 and CHIR, where combinatorial treatment leads to the emergence of new cell populations (Figure S10D,E).

To further assess to what degree information learned in one dataset could help make predictions on other datasets, we next focused on pathway modulators that were used in multiple datasets and systematically withheld all conditions including them from one dataset at a time during training. We compared predictions against a baseline distribution comprising all training conditions from the respective dataset, to evaluate the prediction of effects beyond the protocol differences between datasets. CellFlow largely outperformed this baseline in two datasets (1.7x mean improvement in energy distance) but showed limited predictive power in the Azbukina et al. dataset (Figure S10F,G). We hypothesize that this stems from the exclusive use of higher-order modulator combinations in this dataset (Figure S10H,I), making it challenging to extrapolate from the simpler treatment conditions present in the other datasets. Generally, we conclude that CellFlow enables prediction of cellular compositions arising from new combinatorial treatments in organoid protocols, and shows promising cross-dataset transfer ability for lower-order morphogen combinations.

To systematically explore the space of untested organoid protocols, we used CellFlow to perform a proof-of-principle virtual organoid protocol screen. Based on our previous observations of reduced prediction accuracy for higher-order combinations, we restricted the analysis to single, double, and triple modulator treatments. We focused on 14 pathway modulators that were well-represented across datasets (≥4 conditions) at their highest observed concentration. We then composed protocol conditions by applying each modulator in one of 6 time windows during development (Figure 6F). We trained CellFlow on all available conditions from the three datasets to obtain predictions for all composed protocols. Altogether, our systematic *in silico* screen generated cellular compositions for more than 23,000 protocols, comprising over 70 million cells. We annotated all generated cells through transfer of region, cell type and cluster labels from the primary reference atlas. This revealed that generated organoids were predominantly composed of progenitor and neuron populations with regional identities spanning all major brain regions (Figure 6G and Figure S10J). Overall, the base protocol had a strong impact on brain region and cell type composition. We note that generated organoids corresponding to one base protocol (Sanchis-Calleja et al.) also contained neural crest populations, consistent with observations from the original experimental study where specific morphogen combinations induced neural crest ^47^. These differences between base protocols are expected, as protocol-specific experimental parameters can strongly influence organoid development ^45^. To assess the impact of combinatorial modulator effects across base protocols, we analyzed their influence on brain region composition using a regularized linear model (Methods). We found that both individual modulators and specific combinations showed consistent relationships with specific brain regions across protocols (Figure S10K). For example, retinoic acid (RA) treatment was positively associated with pons and medulla cell identities, while the combination of BMP4 and RA promoted cerebellar fates. This combination in isolation was not included in any of the training datasets, but the predicted effect is consistent with known dorsalizing effects of BMP4 and role of RA in hindbrain patterning ^52–55^.

To further characterize all predicted protocols, we computed two complementary metrics. To assess the biological plausibility of predicted cell populations, we quantified the energy distance between each predicted cell cluster with its counterpart in the reference atlas, then computed a weighted mean across clusters for each condition. We quantified the novelty of predictions with respect to the training data with the energy distance to the closest condition in the training datasets (Figure 6H). Most protocols generated cell populations that closely resembled both primary reference data and existing training conditions, indicating that CellFlow’s predictions largely remained within the bounds of plausible cellular states. A subset of protocols exhibited both high realism (by reference similarity) and novelty compared to the training data. These predictions spanned a range of predicted cellular compositions and naturally clustered into three major categories based on regional identity: mid/hindbrain, ventral forebrain, and dorsal forebrain (Figure S10L). We examined an example from each of these categories as UMAP embeddings, showing trajectories from progenitor to neuron states with distinct brain regional identities that are specific to the protocol composition (Figure 6I). Together, these results demonstrate that CellFlow can generate cell type distributions that resemble organoid and primary cell states and make predictions for untested combinations of morphogen treatments and timing regimens. As a result, CellFlow can help explore and iterate organoid protocol design.

## Discussion

Computational approaches that can learn from high-content phenotypic screens have the potential to accelerate scientific discovery. Our work demonstrates that CellFlow can effectively learn from existing phenotypic screens to predict cellular responses to uncharacterized conditions. CellFlow builds upon flow matching and learns a functional embedding space from experimental variables to encode sophisticated experimental designs. We showed that CellFlow accurately predicts complex phenotypic distributions across diverse biological contexts. We observed a clear scaling law with respect to the number of training conditions, demonstrated by the relationship between prediction accuracy and the number of observed cytokine treatments in the PBMC analysis. This scalable performance indicates that as more perturbation data becomes available, such models will become increasingly accurate and generalizable ^5^.

A key strength of CellFlow is its ability to model heterogeneous cellular distributions that arise from perturbations in developmentally dynamic systems. Other perturbation modeling approaches often struggle to capture the heterogeneous outcomes of fate programming experiments, where interventions can generate diverse mixtures of cell states rather than uniform shifts. Our results demonstrate CellFlow’s capacity to model developmental effects of genetic perturbations across entire zebrafish embryos and predict the emergence of new cell populations in neuron fate engineering as well as organoid cell type composition from various protocol configurations. These applications highlight the importance of modeling distributional perturbation effects, especially in all contexts where cellular heterogeneity is integral to the studied biological system.

Our analyses highlight that out-of-distribution prediction is still a very challenging task. In particular, the effectiveness of predicting responses to entirely unseen perturbations strongly depends on the quality of condition embeddings and their ability to capture functional properties. While we leveraged existing embeddings like ESM2 for proteins and molecular fingerprints for small molecules, these representations may not optimally encode the functional roles relevant to cellular responses. This limitation was evident in our results for completely unseen cytokines or gene knockouts. Finding more powerful representations of perturbed molecular entities that better capture their functional roles could significantly improve extrapolation to unseen contexts. In this context, CellFlow’s trained condition embedding space offers a promising avenue to learn mappings between structure and function.

The modular architecture of CellFlow provides flexibility for extension and improvement across multiple aspects. For cell representation, we explored both PCA and VAE embeddings and found that they present a trade-off between reconstruction capability and flexibility. Future work could explore alternative cell embedding approaches, such as those derived from singlecell foundation models ^42,56^ or disentangled representations ^57^. Moreover, modifications to the flow matching module provide promising avenues for improving CellFlow, for example by directly generating count data ^58^. Stochastic extensions ^59,60^ could enable modeling of the inherent randomness in cellular responses, allowing for better uncertainty estimation on single-cell level, while stochastic parameterization of the encoder module would allow for uncertainty estimation on a distributional level. Furthermore, fine tuning approaches, which are an established part of the training pipeline in computer vision ^61^, could enhance the ability to capture subtle transcriptomic differences. All of these technical improvements could be readily integrated into CellFlow’s modular framework, enhancing its capabilities without requiring major modifications of the overall architecture.

Altogether, CellFlow presents a significant advance in computational modeling of perturbed single-cell phenotypes across diverse biological contexts. By addressing the limitations of existing approaches and providing a modular architecture for ongoing improvement, CellFlow has the potential to accelerate scientific discovery through systematic learning from phenotypic screens. One particularly exciting perspective is the integration of computational modeling with experimental design. Screens can be tailored to exploit the strengths of CellFlow in predicting combinatorial perturbations through sparse experimental designs, and predictions can then point to informative follow-up experiments. This lab-in-the-loop approach could be particularly valuable for exploring complex biological systems, where the space of possible experiments is large but resources are limited. As bigger and more diverse perturbation data sets become available, we anticipate that CellFlow will be a valuable tool to explore the vast space of cellular phenotypes.

## ACKNOWLEDGEMENTS

We thank Larsen Vornholz for input on the PBMC dataset and Lauren Saunders for valuable input on the preprocessing of the zebrafish data. We thank Leon Hetzel for assistance on running chemCPA on the sciPlex3 dataset. This work was funded by the European Union (DeepCell ERC, 101054957), in part by Schmidt Futures for the Cell Annotation Platform, supported by the Helmholtz Association’s Initiative and Networking Fund through CausalCellDynamics (grant #Interlabs-0029). It was supported by the German Federal Ministry of Education and Research (BMBF) under grant no. 031L0289A and 01EE2303E. Moreover, this project has received funding from the European Union’s ERA_PerMed-NET Transcan-3 under grant agreement no. 01KT2308.

## CONTRIBUTIONS

D.K. and J.S.F. conceived the project, with input from A.R., B.T., G.C., and F.J.T.; D.K., J.S.F, and D.B. implemented CellFlow with contributions from G.H.; D.K. benchmarked and conducted analyses of the PBMC dataset and zebrafish dataset with input from J.S.F and A.T.L.; D.K. applied CellFlow to the sciPlex3 and combosciplex dataset, with support from A.S. and M.G. for the benchmark of competing methods; S.B. and L.D. benchmarked CellFlow and competing methods for predicting the effect of combinations of genetic perturbations with input from D.K.; L.Z. and A.P. conducted analyses for the dataset capturing genetic knockouts on cell lines treated with cytokines, with input from D.K. and G.H.; D.B. evaluated CellFlow for neuron fate engineering with support from J.S.F.; J.S.F. evaluated CellFlow on organoid morphogen screens with input from D.B.; J.S.F. performed and analyzed the virtual organoid protocol screen; H.-C.L., N.A., and F.S.C generated the unpublished iNeurons and organoid morphogen screen data; D.K., J.S.F., G.C., and F.J.T. wrote the manuscript with contributions from all authors; D.K., T.U., J.S.F. and D.B. wrote the methods; G.C. and F.J.T. supervised the project; All authors read and approved the final manuscript.

## DATA AVAILABILITY

We make processed versions of the following datasets available via the dataset module of our software package: ^10^milliion PBMCs ^8^, developing zebrafish ^10^, genetic knockouts on cytokine-treated cell lines, and iNeuron ^13^. The sciPlex^34^ and combosciplex ^6^ datasets are available via pertpy ^62^. The 4i data is available via the GitHub repository of CellOT ^17^, and the transcription factor overexpression dataset via the GitHub repository of biolord ^19^. The Amin organoid dataset is available on GEO (GSE233574). The other organoid datasets were obtained from the authors and will be made available through the original studies ^47,49^.

## CODE AVAILABILITY

The CellFlow software package is available at cellflow.readthedocs.io/en/latest/ including documentation and tutorials. The source code is available on github.com/theislab/CellFlow, and code to reproduce our analysis is available at github.com/theislab/cellflow_reproducibility.

## Supplementary Figures

**Figure S1.**
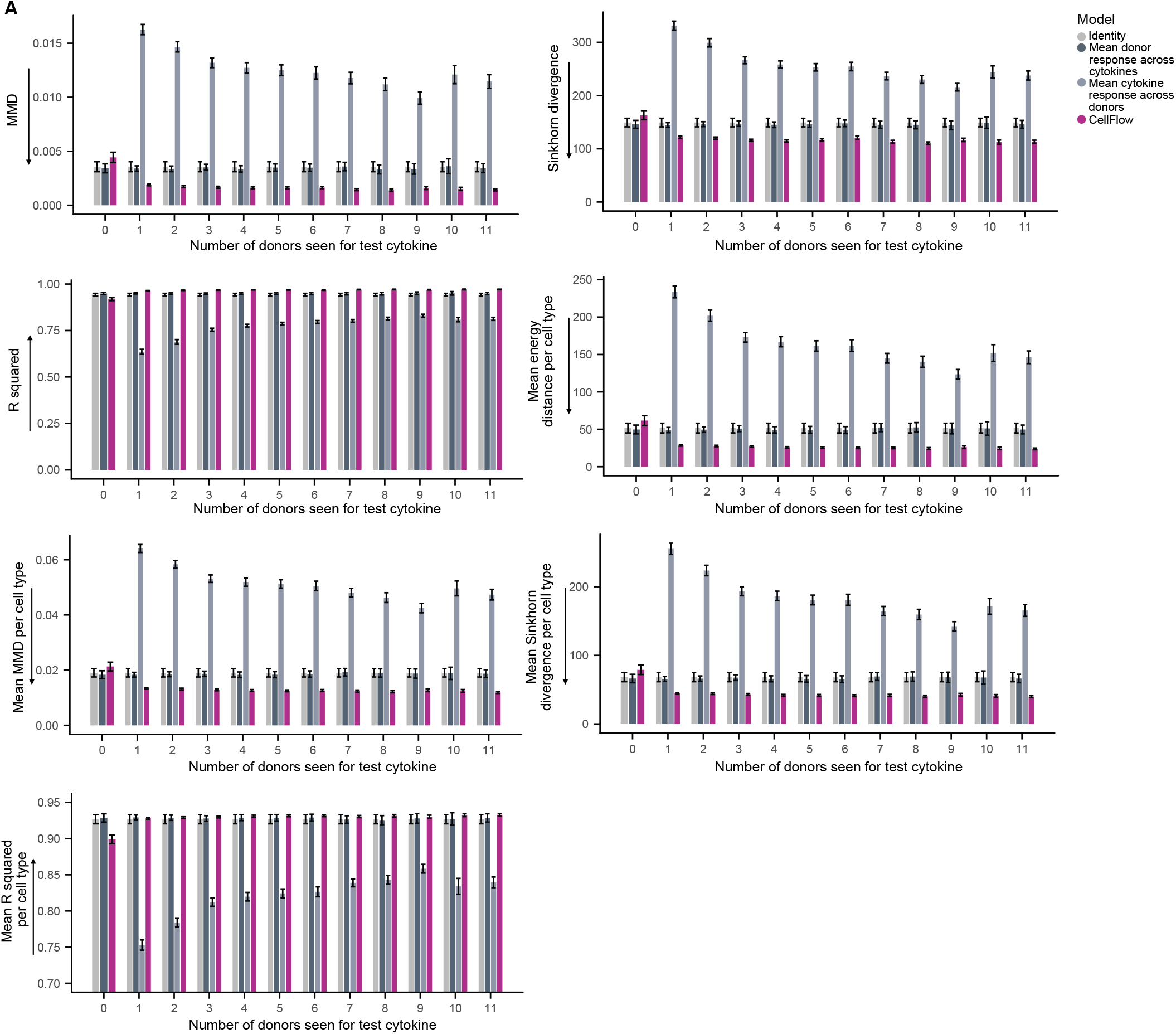
CellFlow is superior to baseline models with respect to different metrics for predicting the effect of a new cytokine. **(A)** Performance metrics MMD (Maximum mean discrepancy) and Sinkhorn divergence in latent space, and R squared of true and predicted mean normalized gene expression for CellFlow and baseline methods reported as mean and standard error across different test cytokines and different sets of donors of size *k* = 0, …, 11 which the test cytokine has been seen for. Analogously, performance metrics are reported per cell type, where cell type labels are transferred using onenearest neighbor (Methods).

**Figure S2.**
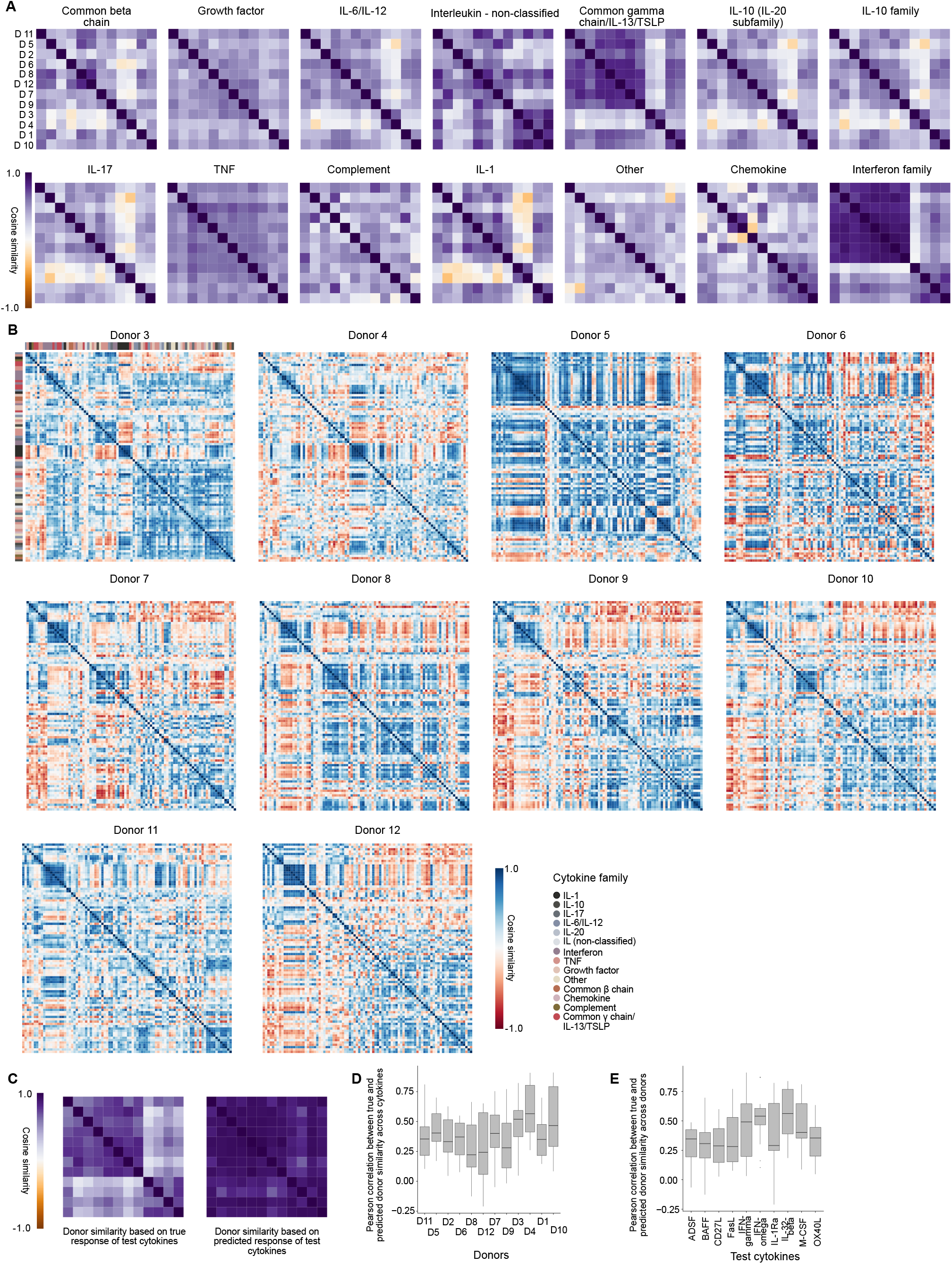
Donor and cytokine similarities of cytokine-treated PBMCs. **(A)** Donor similarities computed from treatment responses of cytokines belonging to one cytokine family (Methods). **(B)** Cytokine similarities computed from treatment responses per donor. **(C)** Donor similarities computed from true cytokine responses based on the ten test cytokines (left), and donor similarities computed from predicted cytokine responses based on the test test cytokines. Predictions were computed from the models including 80 training cytokines for the test donor. **(D)** Pearson correlation between true and predicted donor similarities separated by donor. One data point in a box plot corresponds to one Pearson correlation coefficient between true and predicted donor similarity based on the response of one test cytokine. **(E)** Analogous to (D), but with one data point of the box plot denoting the Pearson correlation coefficient between the true and the predicted donor similarities for one test cytokine.

**Figure S3.**
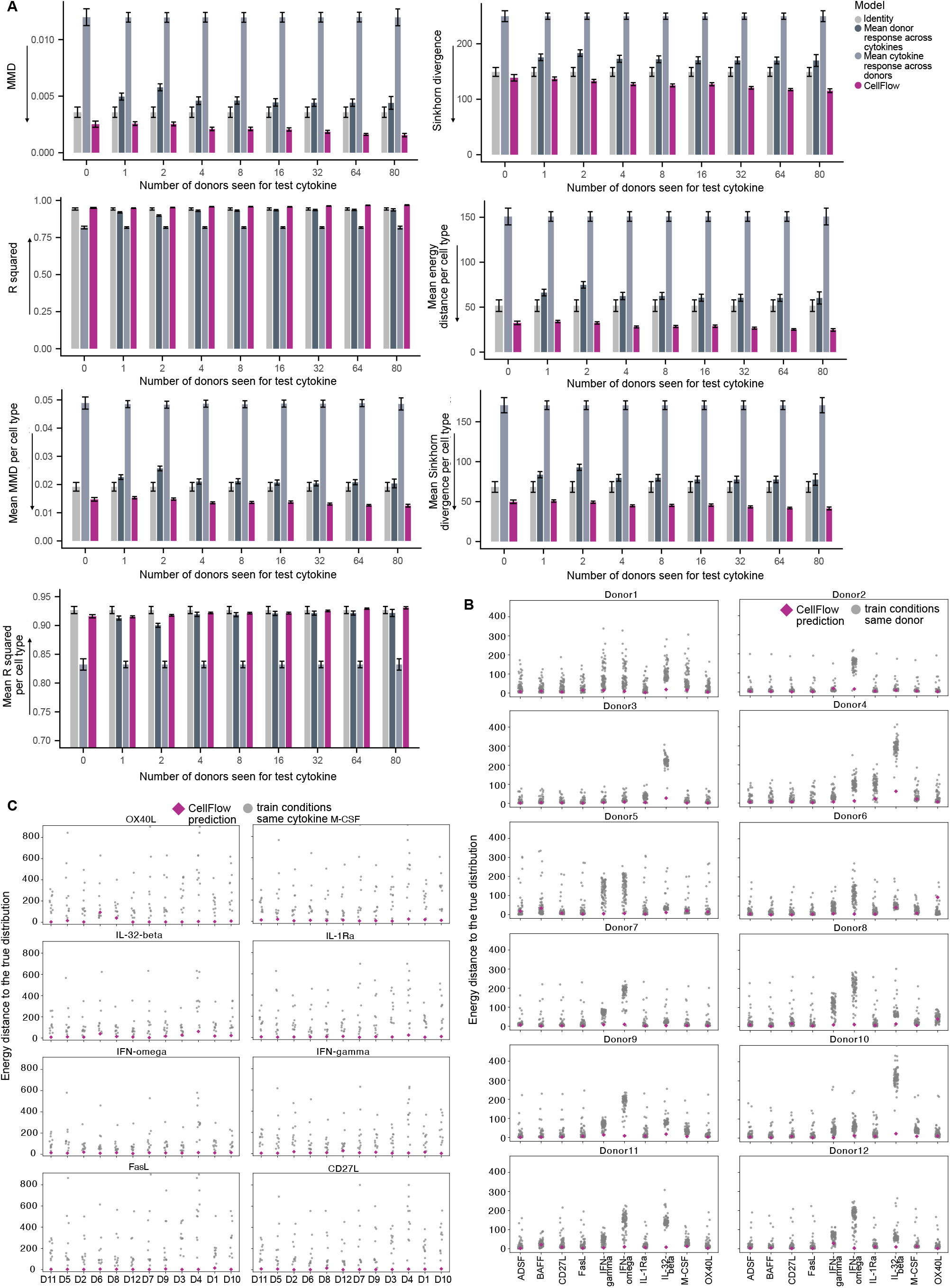
CellFlow out-performs baseline models for predicting the response of a new donor across metrics. **(A)** Performance metrics MMD (Maximum mean discrepancy) and Sinkhorn divergence in latent space, and R squared of true and predicted mean normalized gene expression for CellFlow and baseline methods reported as mean and standard error across different test donors and different sets of cytokines of size *k* which have been seen for the test donor. Analogously, performance metrics are reported per cell type, where cell type labels are transferred using one-nearest neighbor (Methods). **(B)** Comparisons of the distance of CellFlow’s prediction to the ground truth cell distribution (magenta diamond) with distances of training populations of the same donor to the ground truth. CellFlow’s predictions are based on model trainings including 80 cytokines for the test donor. **(C)** Comparisons of the distance of CellFlow’s prediction to the ground truth cell distribution with distances of training populations of the same cytokine to the ground truth.

**Figure S4.**
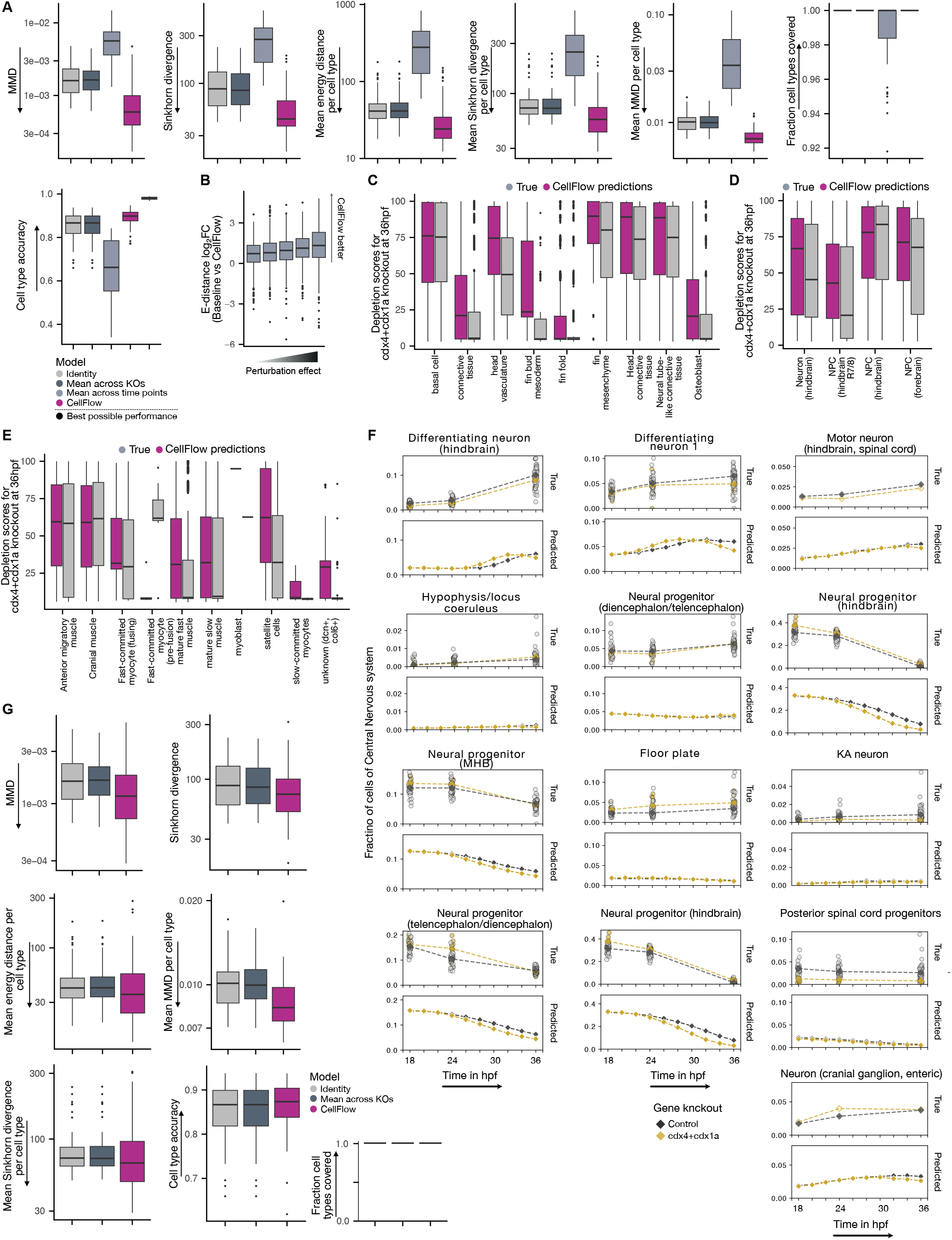
Developmental modeling of perturbed zebrafish embryos. **(A)** Additional metrics for the task of predicting the cellular phenotype of a mutant at an unseen combination of gene knockout and time point (Methods). **(B)** Comparison of CellFlow with the best base line method stratified with respect to perturbation strength (Methods). **(C)** True and predicted depletion scores aggregated to cell type level for the fin. **(D)** Analogous to (c) for neural progenitors. **(E)** Analogous to (C) for cell types of the muscle and connective tissue. **(F)** True and predicted cell type proportions of cdx4+cdx1a mutants at different developmental stages. The training data contains observations of cdx4+cdx1a mutants at time points 18hpf and 36hpf, but not at 24hpf. Observations for other time points are not given in the dataset, and hence only CellFlow’s predictions can be reported. **(G)** Additional metrics for the task of predicting the cellular phenotype of a completely unseen mutation across all time points (Methods).

**Figure S5.**
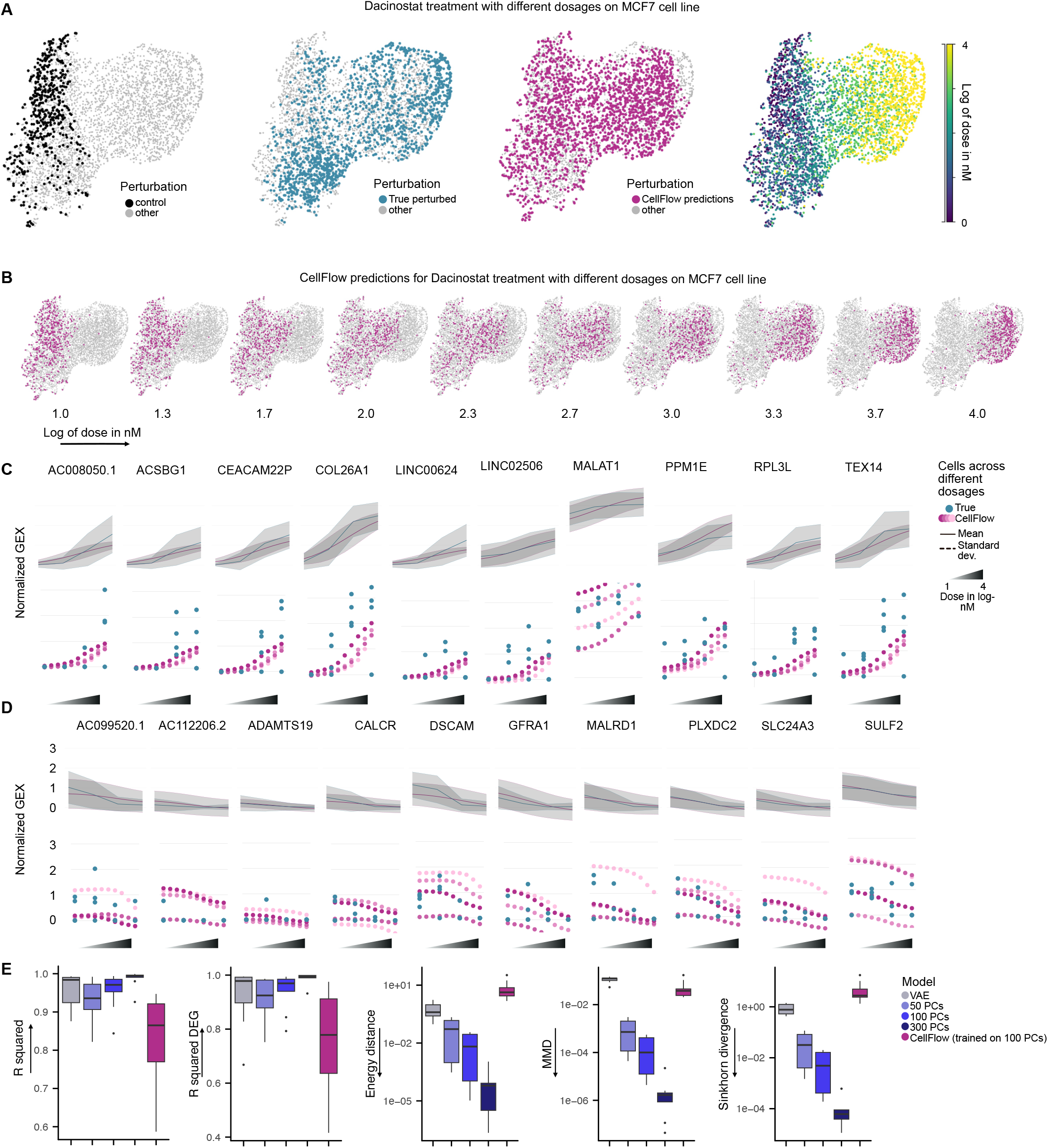
Interpolation of effects across drug dosages. **(A)** UMAP of control cells of MCF7, along with true and predicted Dacinostat-treated cells for dosages 10, 100, 1000, and 10000nM, colored by whether cells are true control, true perturbed, or predicted perturbed cells (from left to right), as well as altogether colored by dose. **(B)** Generated cells for Dacinostat treatment at interpolated dosages. **(C)** Normalized expression of top upregulated genes for Dacinostat treatment of MCF7 with 10000nM. **(D)** Normalized gene expression of top downregulated genes for Dacinostat treatment of MCF7 with 10000nM. **(E)** Upper bound of CellFlow’s performance by encoding-decoding method (Methods) computed by encoding and decoding the true perturbed population, compared with actually achieved performance of CellFlow (trained on 100 PCs).

**Figure S6.**
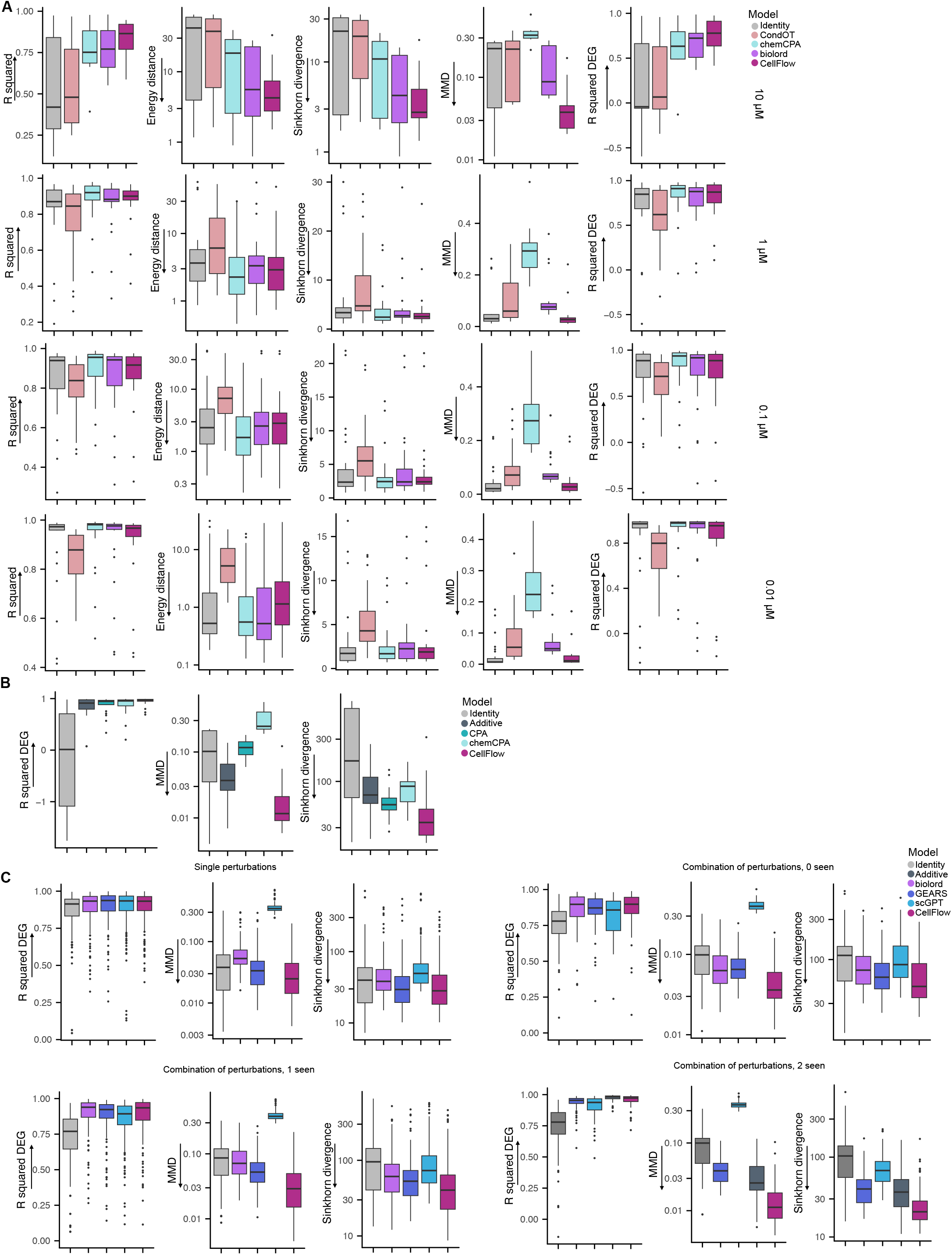
Performance metrics on various perturbation effect prediction tasks. Additional metrics for the task of **(A)** predicting the effect of unseen drugs on the sciPlex3 dataset for different dosages **(B)** predicting the effect of combinations of drugs on the combosciplex dataset and **(C)** predicting the effect of genetic perturbations.

**Figure S7.**
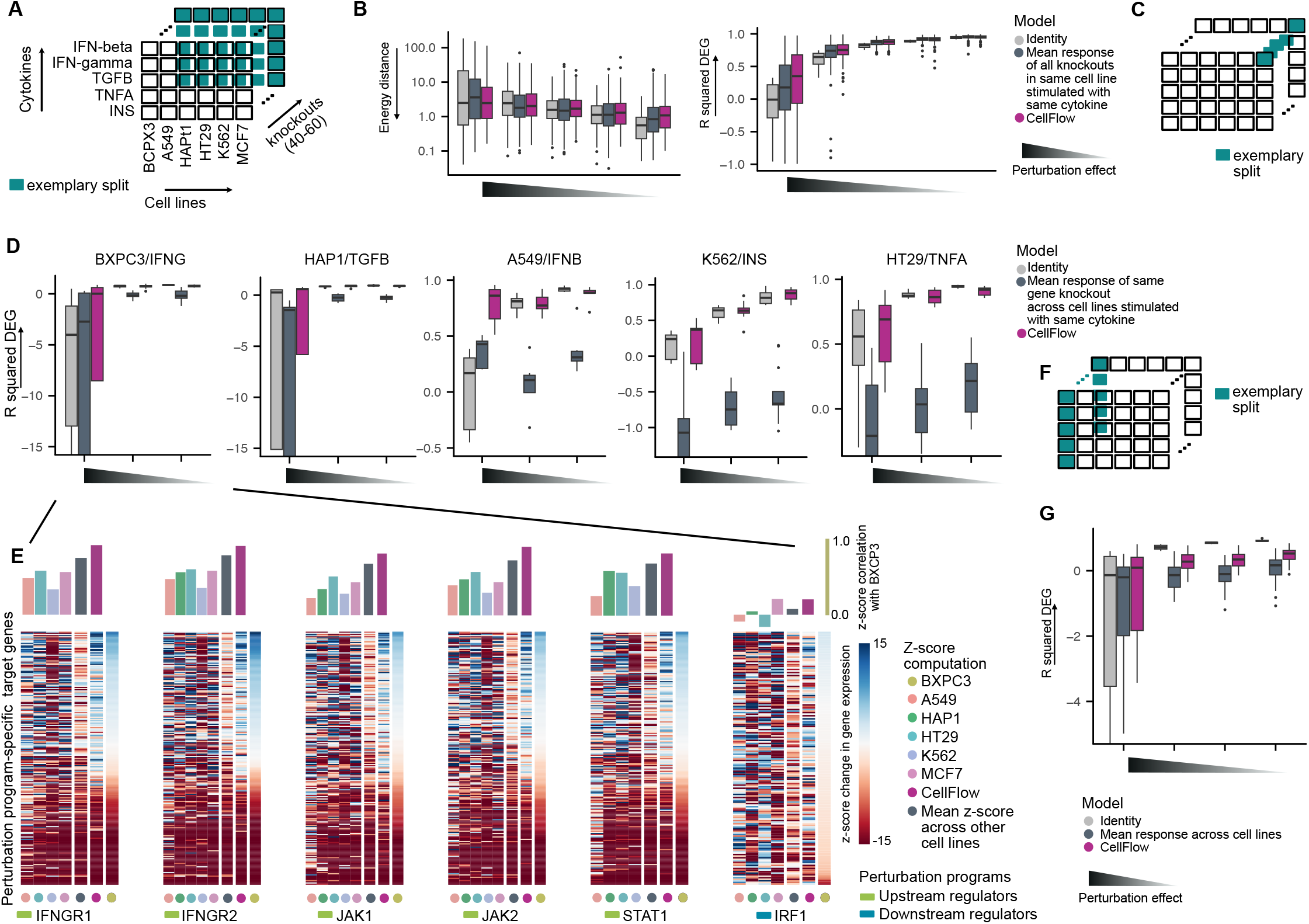
Gene knockout effect prediction across cytokine-stimulated cell lines. **(A)** The dataset contains measurements of gene knockouts which are specific to the cytokine treatment of cell lines ^7^. **(B)** Quantification of the performance of CellFlow for predicting the effect of unseen gene knockouts in cytokine-stimulated cell lines (Methods). Reported are the energy distance in latent space and the R-squared of the condition-specific differentially expressed genes (DEGs) (Methods). Baseline models are the identity, which predicts the perturbation to have no effect, and the mean model, which predicts the perturbation to have the mean effect of the remaining perturbations in the same cytokine-treated cell line. Perturbations are binned into five equally sized groups based on the strength of the perturbation (measured by R squared of DEGs). **(C)** Sketch of the experimental setup of predicting gene knockouts in a new combination of cytokines and cell line, while the cell line and the cytokine have been seen in other combinations in the training data (Methods). **(D)** R squared of normalized gene expression of condition-specific DEGs for each combination of cell line and cytokine which has been held out from the training set. Gene knockouts are evenly split into three groups according to the perturbation strength (measured by R squared of DEGs). **(E)** Z-scores of up- and downregulation of selected genes specific to perturbation programs 7. **(F)** Sketch of the experimental setup of holding out a complete cell line from the training data. **(G)** R squared on DEGs for the task of predicting the effect of all gene knockouts on the BXPC3 cell line for all cytokine treatments for CellFlow and baseline models. For all plots, \**p <* 10^−2^, \*\**p <* 10^−3^, \*\*\**p <* 10^−4^ (unpaired t-test).

**Figure S8.**
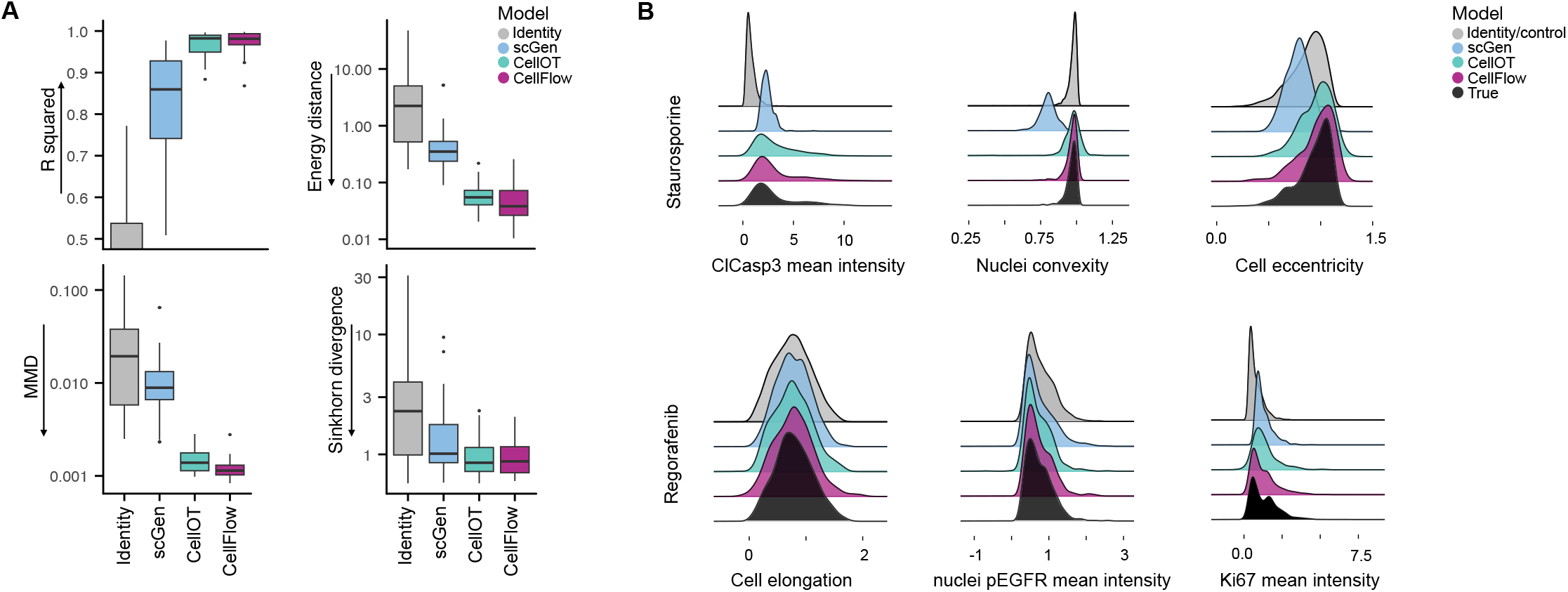
CellFlow predicts the effect of drug perturbations on 4i data. **(A)** Performance metrics for predicting the effect of seen drugs on unseen cells of CellFlow, scGen ^15^, CellOT ^17^, CellFlow, and the identity model, which assumes the drugs to have no effect. The 4i dataset contains readouts for 35 drug treatments on two melanoma cell lines ^17^ (Methods). **(B)** True and predicted marginal distributions of a selected set of molecular and morphological features for two different drug treatments.

**Figure S9.**
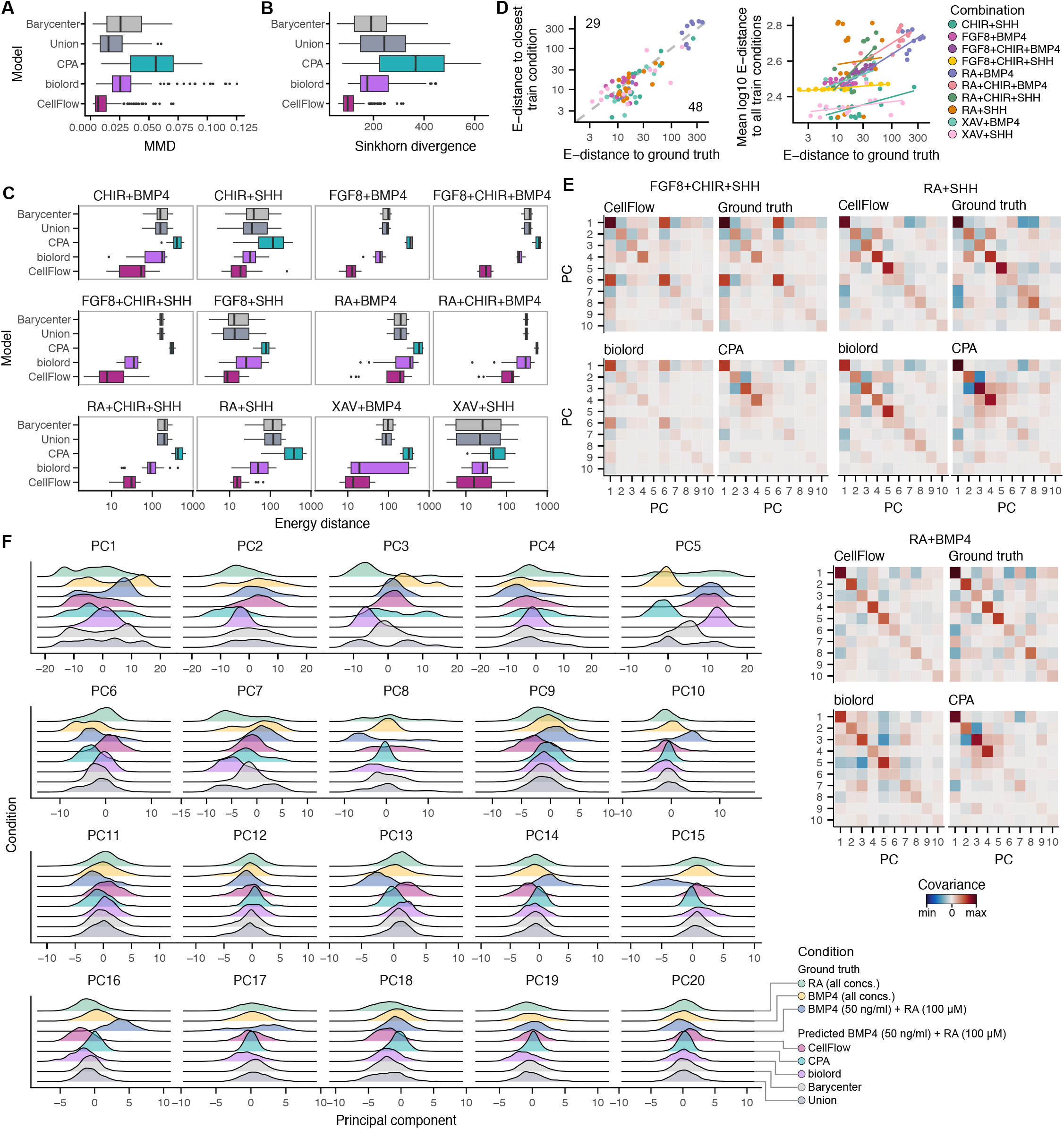
Performance of neuron fate prediction from combinatorial morphogen treatment. **(A and B)** Boxplot showing maximum mean discrepancy (MMD) **(A)** Sinkhorn divergence **(B)** and between predicted and true cell distributions for baselines, other established methods and CellFlow. **(C)** Boxplots showing Energy distance between predicted and true cell distributions for individual morphogen combinations. **(D)** Scatterplots showing the relationship between model performance and energy distance to the closest training condition (left) and mean log_10_(energy distance) across all training conditions (right). Lines represent the linear fits per combination. **(E)** Heatmap showing covariance between the first ten principal components of different conditions for the ground truth, as well as CellFlow, biolord and CPA predictions. **(F)** Density plots showing marginal distributions over all principal components for the true BMP4, RA and BMP4+RA condition as well as the BMP4+RA condition predicted by CellFlow, CPA, biolord models and the barycenter and union baselines.

**Figure S10.**
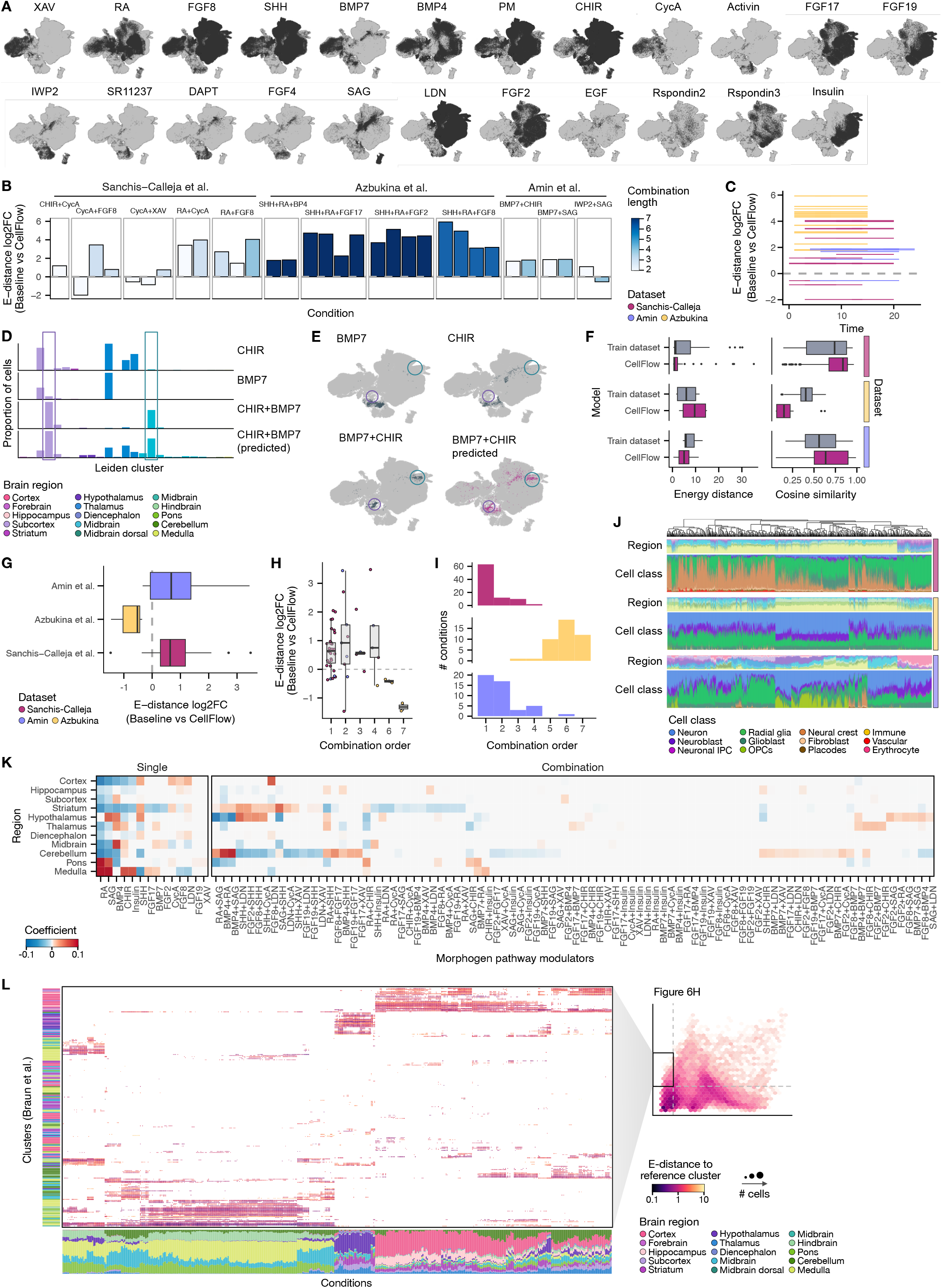
Prediction of organoid cell type composition through digital protocol encodings. **(A)** UMAP embeddings of three combined morphogen screen datasets colored by usage of pathway modulators in screen conditions. All screen conditions using the respective morphogen are colors in dark grey. **(B)** Bar plot showing performance of organoid composition prediction from combinatorial morphogen treatment for all hold-out splits and conditions. Energy distance log2 fold change was computed between CellFlow predictions and the union baseline. **(C)** Treatment time windows and performance for all evaluation conditions. **(D)** Bar plot showing the leiden cluster proportions for the true BMP7, CHIR and BMP7+CHIR condition as well as the BMP7+CHIR condition predicted by CellFlow. **(E)** UMAP embedding showing the cells belonging to the true BMP7, CHIR and BMP7+CHIR conditions (left), BMP4+RA cells predicted by CellFlow projected onto the UMAP embedding (middle). **(F-H)** Performance of predicting the effects of pathway modulators that were held out in one dataset at a time. The baseline comprises all training conditions from the respective dataset to evaluate prediction performance beyond the dataset label. **(F)** Box plots showing the performance in terms of energy distance and cosine similarity of predicted leiden clusters. **(G and H)** Box plot showing log2 fold change of energy distances between CellFlow predictions and baseline split by dataset **(G)** and combination length **(H). (I)** Histogram showing the distribution of combination lengths (how many morphogens were combined) across all conditions in the three datasets. **(J)** Bar pot showing brain region and cell class composition of all predictions from the virtual protocol screen. **(K)** Heatmap showing the coefficients of a L1-regularized linear model to measure the association between pathway modulator treatment and brain region composition based on the virtual protocol screen. **(L)** Dot plot showing the energy distance between predicted cell clusters and corresponding primary reference clusters ^50^ for selected protocols from the virtual protocol screen (energy distance to training data > 1, mean energy distance to reference < 2).

## CellFlow methods

## 1 CellFlow

Given a population of control cells and an experimental intervention, CellFlow aims to generate a perturbed population of cells. We represent the phenotypes of both control (source) and perturbed (target) cells as vectors in a multidimensional embedding space created from principal component analysis (PCA) or variational autoencoders (VAEs). To generate a prediction, CellFlow takes into account all variables defining an experimental intervention and embeds them to a single representation vector, which then guides the transformation of control cells into perturbed cells.

This transformation is governed by a neural ordinary differential equation ^1^ (neural ODE), which is parameterized by a time-dependent neural vector field. The neural vector field is a feedforward neural network that takes as inputs encoded experimental conditions, time points at which the vector field should be evaluated, and interpolated cell phenotype vectors at those time points. Here, time serves as an abstract way to represent a transition from the initial (*t* = 0) to the perturbed (*t* = 1) cellular states. The network outputs velocity vectors defining the flow between these states, and the predicted phenotypes of perturbed cells are obtained by integrating the vector field from *t* = 0, where cell phenotypes are those of the given source cells, to *t* = 1 using a numerical ODE solver.

We train the neural vector field using flow matching and optimal transport. Flow matching ^2^ is a simulation-free way of training neural vector fields, enabling fast and stable optimization. Optimal Transport ^3,4^ (OT) aligns control and perturbed cells by minimizing the total displacement cost, and has been shown to accelerate and improve flow matching training ^5–7^. At each training iteration, a batch of source cells, an experimental condition (e.g. drug or gene knockout), and a batch of target cells corresponding to this condition are randomly sampled. The neural vector field is evaluated at time points which are sampled uniformly from [0, 1]. Subsequently, source and target cells from the sampled batches are paired according to the optimal transport solution for straightening paths of the neural vector field which connects control and perturbed cells. The encoder for experimental conditions and the neural vector field are then optimized end-to-end, thus implicitly learning a functional representation of the condition.

In the following, we elaborate on the most relevant concepts underlying CellFlow.

### 1.1 One formulation to unify the setup of a perturbation experiment

To cover a wide range of perturbation experiments, we formulated a generic setup of phenotypic screens (Figure 1c). We partition the set of variables defining a perturbation experiment into three groups, namely perturbations 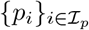, perturbation covariates 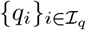, and sample covariates 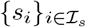, with ℐ_p_, ℐ_q_, and ℐ_s_ denoting index sets. Perturbations include all interventions like drug treatments, gene knockouts, and cytokine treatments, while perturbation covariates add complementary information about the perturbation, e.g. the dosages of drugs or the timing of the intervention. Perturbations (and thus perturbation covariates) can be combined together, for instance, multiple drugs can be given. In contrast, we define sample covariates to capture the cellular state independent of the perturbation. Examples are categorizations of cells into cell lines, tissues, donors, or batches. Sample covariates are not combinable; for instance, a cell cannot belong to multiple batches at the same time.

Thus, a condition **c** a 3-tuple defining the intervention and cellular 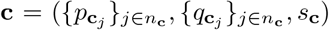. Here, *n*_**c**_ = {1, …, *n*}, *n* ∈ ℕ denotes the number of perturbations (and thus perturbation covariates) of condition **c**. Moreover, condition **c** ∈ 𝒞 is uniquely tied to a pair of source and target distribution (*µ*_**c**_, *ν*_**c**_).

### 1.2 Learning a single representation from all perturbation factors

To learn a representation of the above mentioned factors defining a perturbation experiment, all factors have to be made machine-readable, and subsequently aggregated in a permutation-invariant manner. While there are multiple ways to represent any perturbation, and we leave a systematic evaluation to future work, we employ embeddings from ESM2^8^ (Evolutionary Scale Modeling) to represent cytokines and genetic perturbations, while we leverage molecular fingerprints for drugs ^9,10^. If a representation is not available for the applied treatments, they can be onehot encoded, which only allows predictions of their unseen combinations but not of unseen individual perturbations. As perturbation covariates are typically scalar values (i.e. time of treatment, dosage of treatment), we transform the values to a near-linear scale and neural network-typical range (e.g. applying log-transformation to dosages). However, a user can provide any representation for encoding perturbation covariates. Analogously to perturbations, there is no single representation for all the possible sample covariates. We typically leverage embeddings of the Cancer Cell Line Encyclopedia ^11^ to embed cancer cell lines. For representing patients, we leverage statistics of donor-specific control populations. When learning across datasets, we encode those using one-hot encoding. We refer to an instance defined by perturbations, perturbation covariates, and sample covariates as a condition, and thus aim to learn a representation of conditions which guide the flow from control to perturbed populations.

CellFlow processes the representations of perturbations, perturbation covariates, and sample covariates using modules consisting of feed-forward neural networks or self-attention layers. To obtain a single representation of all cell’s covariates, all of the treatments that make up a single condition are aggregated using a permutation invariant pooling operation. To this end, CellFlow employs permutation-invariant attention mechanisms ^12–14^ or deep sets ^15^ through a feature-wise mean. The resulting condition vector is then further processed through feed-forward neural networks or self-attention modules to obtain a final vector representation of the condition.

### 1.3 Optimal Transport for pairing cells

Pairing samples from the source and the target distribution with OT has proved to improve the generative performance of flow matching models as it straightens paths between source and target populations ^5–7^. Given the complexity and heterogeneity of cellular distributions, mapping control cells to distributionally close perturbed cells is thus a promising direction to improve the generative capabilities of CellFlow.

In the following, we provide background on how to use optimal transport for pairing cells between a source (control) distribution and a target (perturbed) population, before extending the concept to learning a set of optimal transport couplings which allow to map from multiple source to multiple target populations, i.e. when multiple experimental conditions are present. Thus, for the time being, the goal is to find correspondence between source and target cells from the given samples, without any generalization to unseen cells and without introducing different conditions.

#### Monge Maps

Monge maps are deterministic functions which map a source distribution to a target distribution in the most cost-efficient way. Formally, let 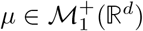 be the source distribution of control cells and 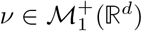 be the target distribution of perturbed cells, where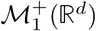 denotes the set of probability distributions on ℝ^*d*^. Our goal is to learn a transport map between these two distributions; that is, to find a map *T* : ℝ^*d*^ → ℝ^*d*^ such that *T ♯ µ* = *ν*. In this case, *T* models the effect of the perturbation: for a control cell **x**~ *µ*, its perturbed version is given by **ŷ** = *T* (**x**) ~ *ν*. To this end, we employ the Optimal Transport (OT) formalism.

Suppose we have a continuous cost function *c* : ℝ^*d*^ × ℝ^*d*^ → ℝ that measures the cost of transporting a control cell **x** ~ *µ* to a perturbed cell **y ~** *ν*. In many applications, this cost is taken to be the squared Euclidean distance between data samples, i.e., *c*(**x, y**) = ∥**x** − **y** ∥ ^2^. We follow this direction in this work. The Monge formulation of OT seeks, among all maps *T* that push *µ* forward onto *ν*, the one that minimizes the expected transport cost:

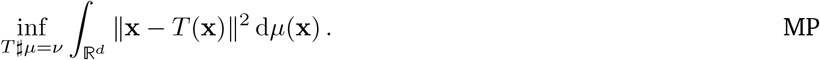

Any solution to Eq. (MP) is referred to as a Monge map between *µ* and *ν*. However, this problem is often difficult to solve in practice because the constraint set can be highly non-convex and may even be empty in discrete settings. Consequently, a relaxed formulation is typically considered instead.

### Kantorovich Formulation

Instead of focusing on transport maps *T*, the formulation introduced by Kantorovich ^16^ considers couplings *π* ∈ *Π*(*µ, ν*). These couplings are probability measures on ℝ^*d*^ × ℝ^*d*^ whose marginal distributions are *µ* and *ν*. Formally,

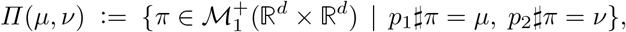

where *p*_1_ : (**x, y**) ↦ **x** and *p*_2_ : (**x, y**) ↦ **y** are the canonical projection maps. Moving from transport maps *T* to couplings *π* replaces a deterministic mapping — where each control cell **x** is mapped to a unique perturbed cell **y** = *T* (**x**) — with a stochastic mapping, where **x** is mapped to a conditional distribution *π*(· | **x**).

Following Eq. (MP), we seek a coupling *π* that minimizes the average displacement cost under *c*:

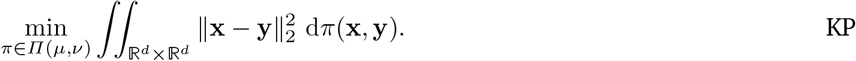

An optimal coupling *π*_OT_ that solves Eq. (KP) always exists. When the Monge map exists, both the Monge and Kantorovich formulations coincide, leading to *π*_OT_ = (Id, *T* ^⋆^)*♯µ*. In other words, *π*_OT_ is a deterministic coupling supported on the graph of *T*. Hence, if (**x, y**) ~ *π*_OT_, then **y** = *T* (**x**).

### Unbalanced Extension

The Kantorovich formulation enforces mass conservation. For instance, this means that every unperturbed cell must be mapped. As a result, factors like apoptosis or asynchronous proliferation cannot be modeled ^7,17^. Unbalanced optimal transport(UOT) ^18,19^ lifts this constraint by penalizing the deviation of *p*_1_*♯π* from *µ* and *p*_2_*♯π* from *ν* via a *ϕ*-divergence *D*_ϕ_. We follow the most common choice *D*_ϕ_ = KL, the KL divergence, defined for positive measures *ρ, γ* ∈ ℳ^+^ (ℝ^*d*^) as

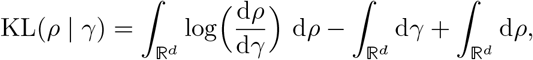

where 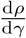 is the relative density of *ρ* with respect to *γ*.

Introducing *λ*_1_, *λ*_2_ *>* 0 to balance how much mass variation is penalized versus transportation, the unbalanced version of the Kantorovich problem seeks a positive measure *π* ∈ ℳ^+^ (ℝ^*d*^ × ℝ^*d*^):

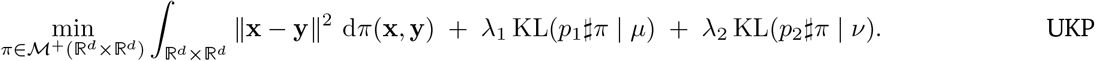

### Entropy-regularized OT

The objective in the Kantorovich problem Eq. (KP) can be smoothed by adding an entropic regularization term. For *ε >* 0, the entropy-regularized Kantorovich problem ^4^ reads

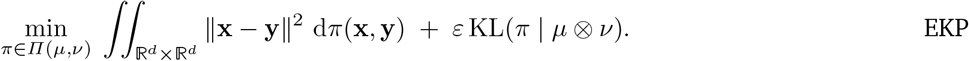

Entropic regularization improves statistical properties by avoiding the so-called OT curse of dimensionality ^20–23^. Moreover, when *µ* and *ν* are discrete, Eq. (EKP) can be solved efficiently by the Sinkhorn algorithm ^4^. This is particularly relevant because, in practice, we only access finite, discrete samples of cells, namely 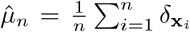 and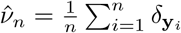, where **x**_1_, …, **x**_n_ and **y**_1_, …, **y**_n_ are i.i.d. samples from *µ* and *ν*, respectively.

In that discrete setting, the entropic Kantorovich Problem Eq. (EKP) translates into an entropy-regularized linear program. By defining the cost matrix **C** = [∥**x**_i_ − **y**_*j*_∥^2^]_*i,j*_ and letting

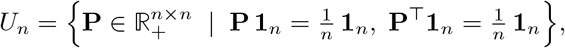

we obtain

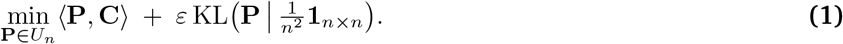

The problem has a dual formulation as an unconstrained, strongly concave maximization:

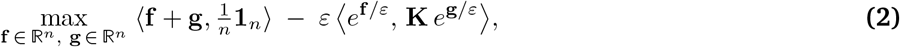

where 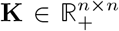 is defined by **K**_i,j_ = exp (−*c*(**x**_i_, **y**_j_)*/ε*), and *e*^**f**^, *e*^**g**^ denote element-wise exponentials. Since the objective is *ε*-strongly concave, an optimal solution **f** ^⋆^, **g**^⋆^ exists and is unique.

Algorithm 1 (the Sinkhorn algorithm) iteratively approximates (**f** ^⋆^, **g**^⋆^). For a matrix **A** = [**A**_*i,j*_]_1≤*i,j*≤*n*_, we define the row-wise *ε*-soft-min operator:

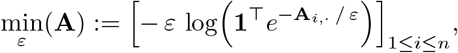

and let ⊕ denote the tensor sum, i.e., **f** ⊕ **g** = [**f**_i_ + **g**_j_]_1≤i,j≤n_. Once **f** ^⋆^, **g**^⋆^ are obtained, the primal-dual relationship

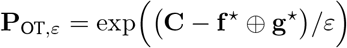

recovers the corresponding optimal coupling. Since each iteration primarily involves matrix-vector multiplications, the time complexity is 𝒪 (*n*^2^), and storing **C** requires 𝒪 (*n*^2^) memory in the most generic case. Yet, when the cost is separable (e.g. the squared Euclidean cost), it is possible to reduce the memory complexity to linear ^24^.

A similar approach applies to the unbalanced problem Eq. (UKP) by adding an entropic penalty. Formally,

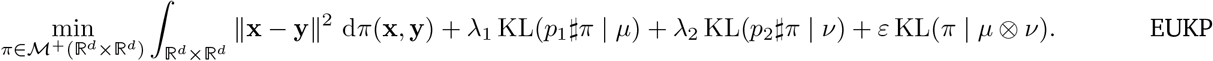

In the discrete case, using similar notations as before, we get

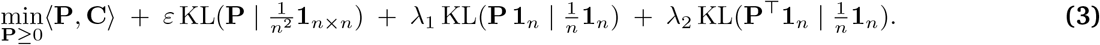

Like the balanced setting Eq. (3), this problem admits a dual formulation as an unconstrained, strongly concave maximization ^25,26^:

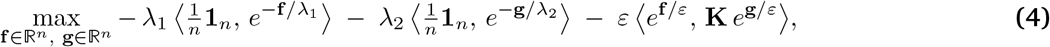

and it has a unique solution (**f** ^⋆^, **g**^⋆^). A variant of the Sinkhorn algorithm, requiring minor modifications, solves this problem. Specifically, we define

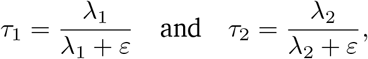

and rescale **f** and **g** by *τ*_1_ and *τ*_2_, respectively. Intuitively, *τ*_1_ and *τ*_2_ govern how closely the row and column sums of **P** must stay near 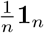. When *λ*_1_, *λ*_2_ → ∞, we recover *τ*_1_ = *τ*_2_ = 1, matching the balanced Sinkhorn algorithm. Algorithm 1 highlights this scheme, which mainly alternates between matrix-vector multiplications in 𝒪 (*n*) time.

After convergence, the primal-dual relationship

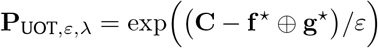

recovers the optimal plan. Unlike the balanced case, row and column sums need not match 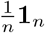 exactly but remain close in a trade-off driven by *λ*_1_, *λ*_2_. In practice, it is often more convenient to tune *τ*_1_ and *τ*_2_ directly, since *τ*_1_, *τ*_2_ ∈ [0, 1]. As a result, we reparametrize the notation in terms of *τ* = (*τ*_1_, *τ*_2_), and denote 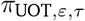 and 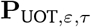 accordingly.

#### Algorithm 1

Sink(**X, Y**, *ε, τ*_1_, *τ*_2_). **Note:** use *τ*_1_ = *τ*_2_ = 1 for balanced Sinkhorn.

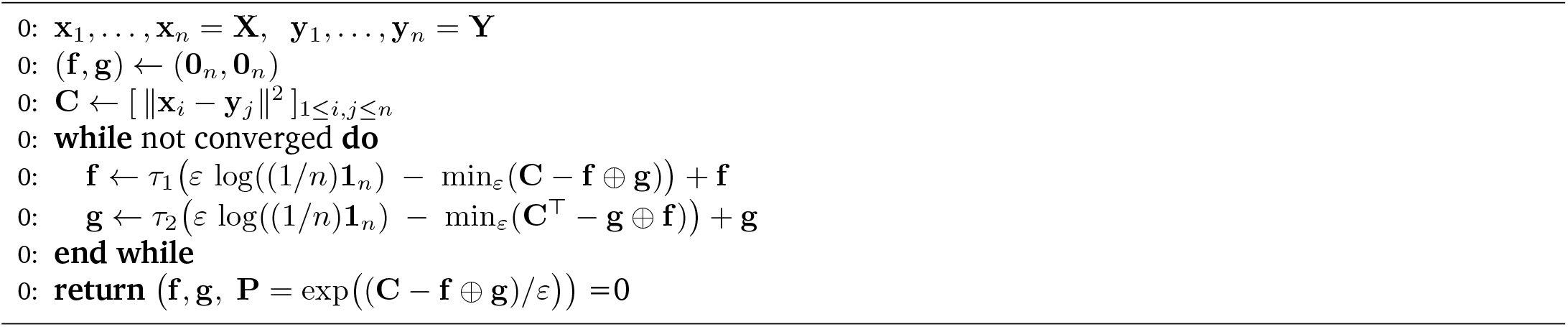

We leverage `ott-jax` for computing OT couplings, and hence it is straightforward to run CellFlow with adaptions like low-rank approximations for particularly large batch sizes and online cost matrix evaluation for reducing memory complexity ^24,27^.

### 1.4 Flow Matching for generating perturbed cell populations

In this section, we introduce flow matching and show how it can be combined with discrete OT techniques to approximate OT maps between distributions of cells. From learning the correspondence between source and target cells, we shift to learning a vector field that describes a transition from one population to the other.

#### Continuous Normalizing Flows (CNFs)

Continuous normalizing flows ^1^ transform a source distribution 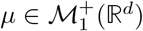 to a target distribution 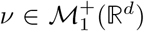. Translated to our setting, the source distribution corresponds to an unperturbed population, while the target distribution corresponds to a perturbed population. Instead of directly parameterizing *T* as a neural network, CNFs make use of the concept of a flow *ϕ*_1_ : ℝ^*d*^ → ℝ^*d*^ which is induced by a velocity field *v*_t_ : ℝ^*d*^ → ℝ^*d*^, namely *ϕ*_t_ : ℝ^*d*^ → ℝ^*d*^ solves:

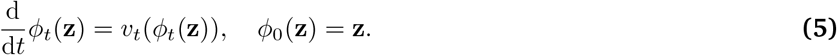

CNFs model the vector field with a neural network *v*_t,θ_, leading to a deep parametric model of the flow, which is trained to match the terminal condition *ϕ*_1,θ_*♯µ* = *ν*.

#### Flow Matching

While the standard way of training CNFs included the typically computationally expensive way of solving a neural ordinary differential equation (ODE), flow matching ^2^ alleviates this constraint, and thus results in a so-called simulation-free technique for training CNFs. It constructs a probability path *p*_t_ that transitions from *p*_0_ ≈ *µ* to *p*_1_ ≈ *ν* by marginalizing individually constructed conditional paths *p*_t_(·| **x, y**) for data samples **x** from *µ* and **y** from *ν*. These conditional paths are used as a ground truth, which the paths generated by the learned vector field should approximate. Formally, we take any coupling *π* ∈ *Π*(*µ, ν*) that has *µ* and *ν* as its marginals—such as the independent coupling *π* = *µ* ⊗ *ν*—and we define for **z** ∈ ℝ^*d*^,

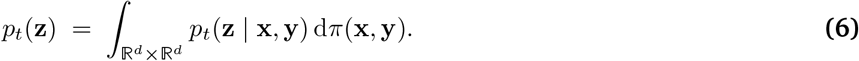

We assume these conditional paths can be generated by conditional velocity fields *u*_t_(·|**x, y**). Then, the marginalized velocity field is given by

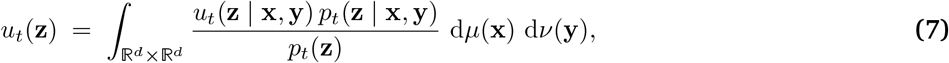

and it can be shown that this *u*_t_(**z**) recovers the marginalized path *p*_t_. In principle, one could then learn *v*_t,θ_(**z**) that approximates *u*_t_(**z**), and thus generates paths that approximate *p*_t_(*z*), by minimizing

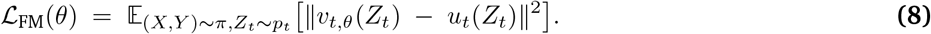

However, this objective is intractable because *u*_t_ depends on *p*_t_. Instead, it was shown ^2^ that it suffices to regress onto the conditional velocity fields and minimize

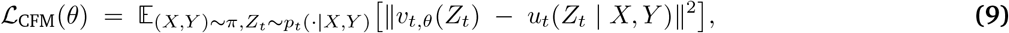

as ∇_θ_ ℒ_CFM_(*θ*) = ∇_θ_ ℒ_FM_(*θ*). Hence, optimizing the tractable loss ℒ_CFM_(*θ*) is equivalent to optimizing ℒ_FM_(*θ*), allowing one to learn *v*_t,θ_ without simulating trajectories by choosing conditional probability paths *p*_t_(· | *X, Y*) and conditional vector fields *u*_t_(*Z*_t_ | *X, Y*) that have a simple form. If this loss is 0, the flow is a push-forward map, i.e., *T* = *ϕ*_1_ satisfies *T ♯µ* ≈ *ν* ^2^ Theorem 1. In other words, this means that the flow induced by the neural velocity field *v*_t,θ_ transports the distribution of control cells to the distribution of perturbed cells.

### Gaussian Probability Paths

A natural choice for the construction of the ground truth probability paths, which we adopt in this work, is to use Gaussian conditional probability paths of the form

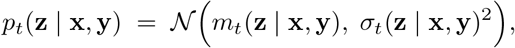

where *µ*_0_(**x, y**) = **x**, *µ*_1_(**x, y**) = **y**, and *σ*_t_(**x, y**) ≥ 0. Assuming that *m*_t_ and *σ*_t_ are differentiable in *t*, the corresponding conditional velocity field

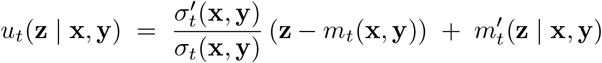

can be shown to induce the above Gaussian path *p*_t_(**z** | **x, y**) via the affine conditional flow

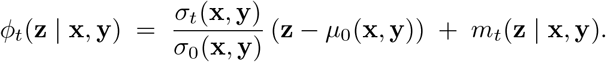

In this work, we always choose a linear interpolation between the data samples as the means along the conditional path, namely

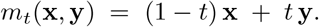

We then consider two different noise schedules:

1. Constant noise:

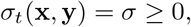

2. Brownian bridge:

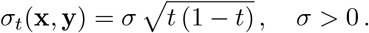

These schedules provide the following conditional velocity fields:

1. Constant noise:

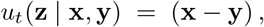
2. Brownian bridge:

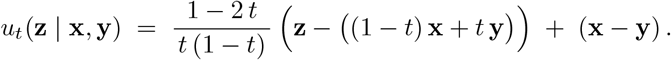

To conclude, we consider (noisy) straight lines as conditional probability paths. On the other hand, while we yield individual straight paths between source and target pairs, the map *ϕ*_1_ induced by the marginalized velocity field *u*_t_ has no reason to be an OT map.

### OT Flow Matching

To promote the flow *ϕ*_1_ to mimic an OT map, Tong et al. ^5^, Pooladian et al. ^6^ propose pairing the samples between which we join the (noisy) straight paths via an (entropic) OT coupling, instead of picking them independently. Specifically, in Eq. (6) we choose 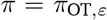 with *ε* ≥ 0, where 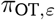 solves Eq. (EKP) when *ε >* 0 or Eq. (KP) when *ε* → 0, between *µ* and *ν*. This yields the OT-CFM loss

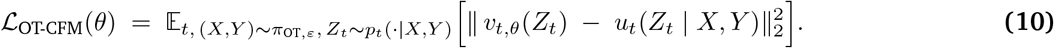

In practice, we do not have direct access to *π*_OT,ε_, so we approximate it from data samples **x**_1_, …, **x**_n_ from *µ* and **y**_1_, …, **y**_n_ from *ν* using the Sinkhorn algorithm (1). When *ε* → 0 and *σ*_t_(**x, y**) = *σ* ≈ 0, *ϕ*_1_?asymptotically approximates a Monge map ^6^ between *µ* and *ν*. On the other hand, for *ε >* 0 and 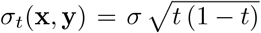, the joint distribution of (*X, ϕ*_1_(*X*)) for *X* ~ *µ* asymptotically approximates that of the solution to Eq. (EKP) between *µ* and *ν* ^28^.

### UOT Flow Matching

Eyring et al. ^7^ extend this idea to the unbalanced setting by pairing samples using the unbalanced OT coupling *π*_UOT,ε,τ_ with *ε* ≥ 0 and *τ* = (*τ*_1_, *τ*_2_) ≥ 0. This coupling solves Eq. (EKP) between *µ* and *ν* when *ε >* 0 or Eq. (KP) when *ε* → 0. This yields the UOT-CFM loss

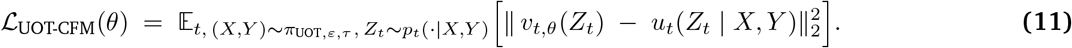

Here, *π*_UOT,ε_ is approximated from samples of *µ* and *ν* using the Sinkhorn algorithm 1. In the limit *ε* → 0 with *σ*_t_(**x, y**) = *σ* ≈ 0, the map *ϕ*_1_ asymptotically approximates a Monge map between the re-weighted versions 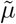 and 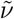 of *µ* and *ν*, which are the marginals of *π*_UOT,ε_. On the other hand, for *ε >* 0 and 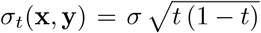, the joint distribution induced by *ϕ*_1_ asymptotically approximates that of the solution to Eq. (EKP) between 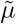 and 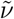. In short, UOT-FM can be regarded as OT-FM benefiting from an automatic (and costless) “pre-conditioning” step on the source and target distributions, thereby simplifying transportation.

### 1.5 The CellFlow algorithm

Having introduced optimal transport couplings and flow matching for a single source distribution *µ* and a single target distribution *ν*, we now extend the setting to multiple pairs of source and target distributions {(*µ*_**c**_, *ν*_**c**_)} _**c**∈𝒞_. We note that this includes the setting of having only one source distribution *µ*, but multiple target distributions *ν*, i.e.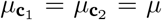, but e.g. 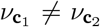. As introduced above, we tie a pair of source and target distribution (*µ*_**c**_, *ν*_**c**_) with variables defining the experimental intervention 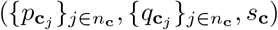 defining the transformation from *µ* to *ν*. We provide the information about the experimental interventions as inputs to the learned vector field, so that, upon training, it could be evaluated given a new experimental condition.

Putting everything together, we can now derive the CellFlow loss functions. We detail the construction of these objectives below, explaining how model choices influence them.

1. First, we set a distribution *ρ* over conditions **c** ∈ 𝒞 which we choose to be uniform, i.e. 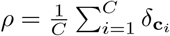.
2. We then embed the perturbation variables 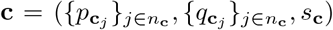 via an aggregator scheme A_θ_ into a condition vector 𝒜_*θ*_(**c**).
3. Given the condition **c** ~ *ρ*, we sample batches of cells from *µ*_**c**_ and *ν*_**c**_. For these batches, we find a coupling *π*_**c**_ ∈ *Π*(*µ*_**c**_, *ν*_**c**_) by setting either *π*_**c**_ = *π*_OT,ε,**c**_ or *π*_**c**_ = *π*_UOT,ε,τ,**c**_ and approximating it using Algorithm 1.
4. Next, we specify a (shared among conditions) family of Gaussian conditional probability paths *p*_t_(**z x, y**) = 𝒩 (*m*_t_(**x, y**), *σ*_t_(**x, y**)^2^) that connect coupled data samples **x** ~ *µ*_**c**_ and **y** ~ *ν*_**c**_ from the conditional source and target distributions. These paths are induced by conditional velocity fields *u*_t_(**z x, y**) obtained via *m*_t_(**x, y**) and *σ*_t_(**x, y**).
5. Finally, we utilize conditional probability paths to sample intermediate cell states *z*_t_ ~ *p*_t_(**z** | **x, y**) (denoted by *X*_t_ in Figure 1). Given these samples and the condition vector 𝒜_*θ*_(**c**), we use neural vector field *v*_t,θ_(**z**_t_ | 𝒜_*θ*_(**c**)) to approximate *u*_t_(**z**).

This yields the CellFlow loss

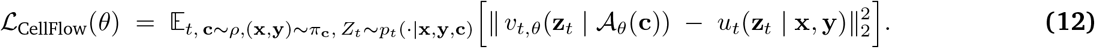

We highlight the CellFlow training procedure in algorithm 2. To consider multiple conditions per batch, we use gradient accumulation over *T*_acc_ steps before updating the parameters.

#### Algorithm 2

CellFlow(*v*_t,θ_, *ρ*, (*µ*_**c**_, *ν*_**c**_)_**c**_, (*π*_**c**_)_**c**_, *m*_t_, *σ*_t_, *n, ε, τ*_1_, *τ*_2_, *T*_acc_).

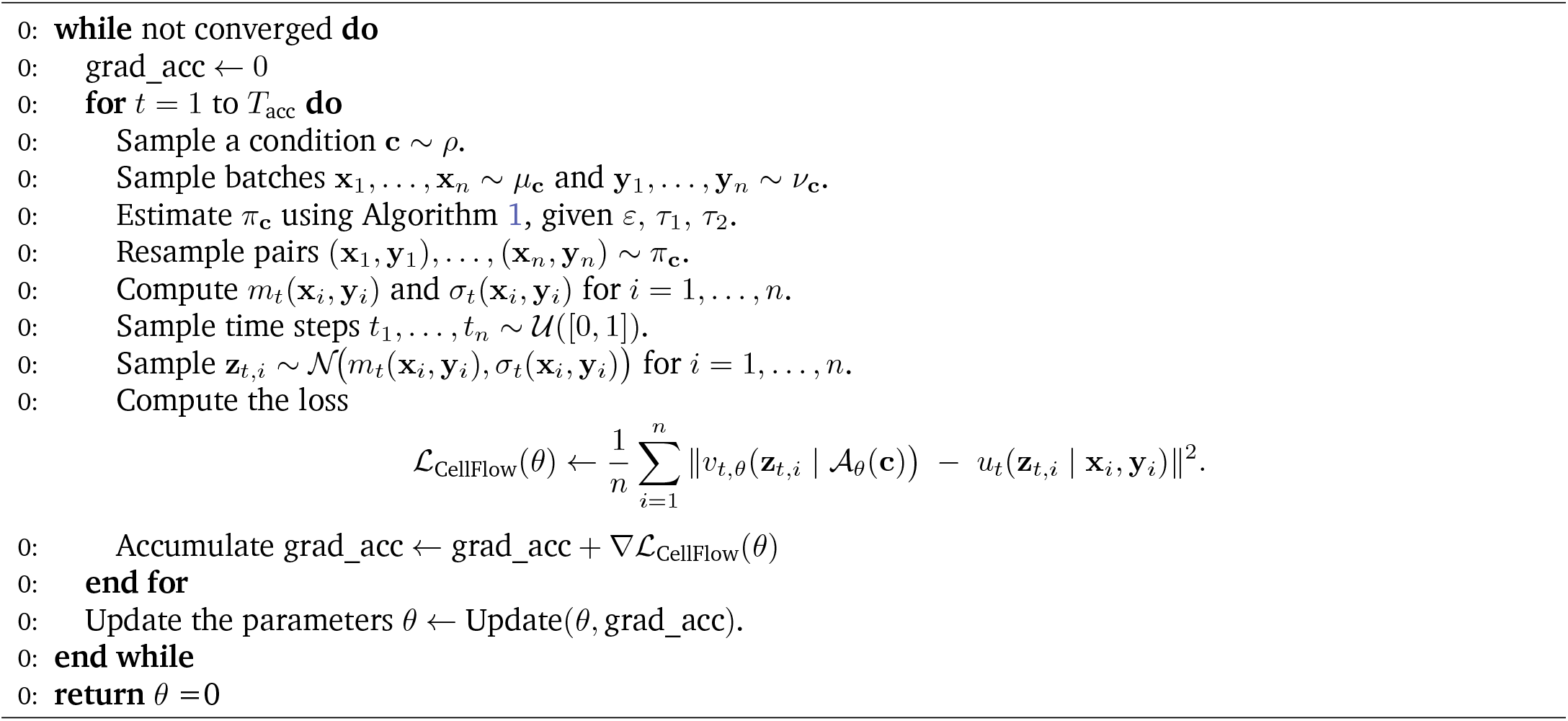

### 1.6 Parameters in CellFlow

While the CellFlow architecture is implemented as and meant to be a flexible framework which can be built upon in future times, we provide an overview of the most relevant parameters in the following, divided into the categories mini-batch coupling, condition encoder, flow matching module, and training parameters.

#### Mini-batch OT coupling

Hyperparameters for the discrete OT coupling are the ones which are commonly used for (linear) optimal transport, and we refer the reader to the documentation of `ott-jax`^24^ for a detailed instruction, and to `moscot` ^30^ for the most relevant ones in the context of single-cell genomics.

It is important to highlight that building upon OT-CFM ^5–7^, we are limited in the choice of the cost to *L*^p^, as otherwise, the coupling information is not necessarily preserved ^**?**^.

The most relevant hyperparameters for the mini-match OT coupling are

- *ε*: entropic regularization ^4^ determining the stochasticity of the coupling. The more ‘complex’ the control and perturbed distribution, the smaller *ε* should be chosen. To prevent dependence on the scale of the data, we always scale the cost matrix by its mean ^30^. Default value: *ε* = 1.0.
- *τ*_a_, *τ*_b_: Left and right unbalancedness parameters between 0 (fully unbalanced) and 1 (fully balanced). The more outliers or distributional undesired biases in the source and target distribution, the smaller the *τ*_a_ and *τ*_b_, respectively. Default value: *τ*_a_ = *τ*_b_ = 1.0.

#### Encoder module

Each perturbation, covariate, and cell line embedding can be further processed by a feedforward neural network (`mlp`) or self-attention module (`self-attention`) before being concatenated, the entirety of which we refer to `layers_before_pool`. These concatenated vectors are then pooled using standard attention (`attention_token`), set attention (`attention_seed`), or mean-pooling/deep sets (`mean`). When the set of vectors to combine is of size one, attention falls back to self-attention, while pooling is equivalent to the identity. The resulting vector is then fed forward through another feedforward neural network (`layers_after_pool`), before it is projected to a vector of size `condition_embedding_dim`, the output of the encoder module. In summary, we have the following parameters together with default values in parantheses

- `Pooling (attention_token)`
- `cond_output_dropout (0.9)`
- `condition_embedding_dim (256)`
- `layers_before_pool (specified for perturbation, perturbation covariate and sample covariate):`
  – `layer_type (mlp)`
  – `dims ([1024, 1024])`
  – `dropout_rate (0.0)`
- `layers_after_pool`
  – `layer_type mlp`
  – `dims ([1024, 1024])`
  – `dropout_rate (0.0)`

**Flow Matching module** The flow matching module transforms the source (control) distribution to a target distribution (perturbed) given a condition. The `time_encoder_dims` is a feed-forward MLP encoding the sinusoidal embedding of dimension `time_freqs` of the time (not experimental time, but time of the neural differential equation). The `hidden_dims` specify the architecture of the MLP processing samples (cells) in the source (control) distribution. The `decoder_dims` define the layers of the MLP processing the concatenated vectors (linearly projected if `linear_projection_before_concatenation`, and layer-normalized if `layer_norm_before_concatenation`) of the condition embedding (from the condition encoder), the time embedding (output of `output_dims`), and the processed cell embedding (output of `hidden_dims`). The noise schedule can be chosen arbitrarily (see 1.4), we recommend adding constant Gaussian noise (`constant_noise`) with standard deviation `flow_noise`, or noise following the noise schedule of a Schroedinger bridge `bridge`.

- `time_freqs (1024)`
- `time_encoder_dims ([2048, 2048, 2048])`
- `time_encoder_dropout (0.0)`
- `hidden_dims ([4096, 4096, 4096])`
- `hidden_dropout (0.0)`
- `linear_projection_before_concatenation (False)`
- `layer_norm_before_concatenation (False)`
- `decoder_dims ([4096, 4096, 4096])`
- `decoder_dropout (0.0)`
- `flow_type (constant_noise)`
- `flow_noise (0.1).`

**Training**

In contrast to most other deep learning methods, samples in one batch (of size `batch_size`) are not independent of each other due to the resampling step according to the OT plan. The number of iterations (`num_iterations`) is a hyperparameter which has to be optimised; due to the relatively costly inference, early stopping is not as cheaply available as in one-step models. `multi_steps` is one of the most relevant parameters. It denotes the number of conditions (not cells) seen between gradient updates. It inherits its name from its implementation, i.e., due to the data structure, we first sample a source distribution (sample covariate), and then corresponding perturbations and perturbation covariates. Given this configuration of conditions, we sample cells from the control and perturbed population, denoting one “step” of the training procedure. As this corresponds to only seeing one data point in the condition encoder module, we increase the number of steps to `multi_steps`, for the condition encoder module not to overfit. In summary, we have the following parameters together with default values

- `batch_size (1024)`
- `num_iterations (500_000)`
- `multi_steps (20)`
- `optimizer (Adam as implemented in optax` ^31^)
- `learning_rate (0.00005)`.

### 1.7 Implementation of CellFlow

CellFlow is implemented in `JAX` ^32^, and thus makes use of its deep learning ecosystem including `flax`, `optax`, and `diffrax`^33^. We make use of `ott-jax`^24^ for pairing cells with optimal transport, and also use their implementations for parts of the flow matching module. For preparing and processing the data, we build upon the scverse ecosystem ^34^ including `anndata` ^35^ and `scanpy` ^36^ as well as its GPU-accelerated counterpart `rapids-singlecell`. For `CellFlow`’s preprocessing module, we build upon `pertpy` ^37^ and `HuggingFace`.

## 2. Metrics

In this section, we introduce several distributional metrics to evaluate how well generative models capture the underlying distribution of perturbed cells. We note that not all of them are actual metrics, but for the sake of readability we refer to them as such.

### 2.1 R squared

R squared (R^2^) between the true and predicted mean gene expression is arguably the most common metric used in the perturbation modeling community ^38–41^. Given true samples 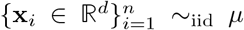 and predicted samples 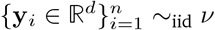, we first compute the empirical means

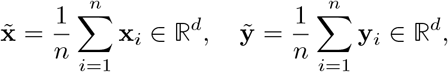

and then define the empirical R^2^ between these two empirical means as

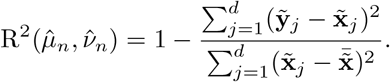

with

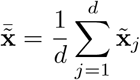

There are two major advantages of the R^2^ metric for evaluation of perturbation predictions. First, it is independent of any hyperparameter choices. Second, it is does not suffer from the curse of dimensionality, i.e. it can be evaluated in very high dimensions without losing expressivity, which distinguished this metric from the remaining ones.

However, R^2^ values must be handled with care. In particular, it only takes into account the mean, i.e. the first moment of a distribution. This entails that it does not distinguish between dimensions of little and dimensions of much variation, weighing each axis equally. Thus, while it might be useful for evaluating predictions on “simple” distributions like cell lines, it is not suitable for evaluating heterogeneous populations.

#### Sample Complexity

Under mild moment conditions and fixed dimension *d*, standard concentration results imply the empirical R^2^ converges to its population counterpart at the parametric rate *O*(*n*^−1/2^).

### 2.2 Maximum Mean Discrepancy

The Maximum Mean Discrepancy (MMD) is a metric for comparing two probability distributions *µ* and *ν* in 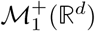.

To define the MMD, we start with a conditionally positive definite kernel, i.e., a function *k* : ℝ^*d*^ × ℝ^*d*^ → R such that, for any points **x**_1_, …, **x**_n_ ∈ R^d^ and real coefficients *a*_1_, …, *a*_n_ satisfying 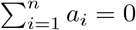, we have

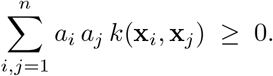

If we remove the requirement 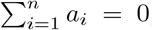, then *k* is called *positive definite*, which is a strictly smaller family contained within the conditionally positive definite kernels.

Given such a kernel *k*, the (squared) MMD between *µ* and *ν* is defined by

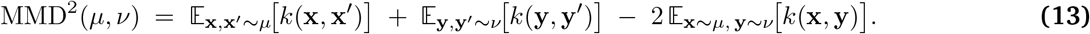

When *k* is characteristic, MMD(*µ, ν*) = 0 if and only if *µ* = *ν*. Thus, MMD truly measures the distributional discrepancy between *µ* and *ν*. As a result, the MMD has been widely used for generative model evaluation and other tasks requiring distributional comparisons.

#### RBF MMD

A popular choice for *k* is the Gaussian radial basis function (RBF) kernel, defined by

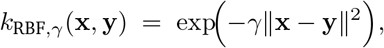

where *γ >* 0 is a bandwidth parameter. This kernel is (conditionally) positive definite on R^d^ and is also characteristic, which ensures that MMD can distinguish between distinct *µ* and *ν*.

By substituting *k*_RBF_ into the MMD definition 13, we obtain

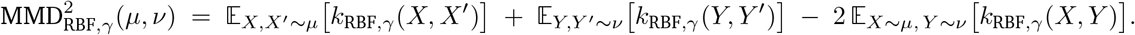

If not stated otherwise, we follow Bunne et al. ^42^ in computing the MMD across different length scales *γ* ∈ {2, 1, 0.5, 0.1, 0.01, 0.005} and reporting its mean.

#### Sample Complexity

In practice, we compute MMD^2^ on empirical distributions 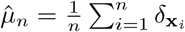 and 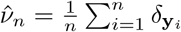leading to

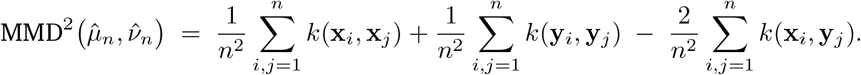

Computationally, this procedure is straightforward because it only involves pairwise kernel evaluations. Additionally, under mild regularity conditions on the kernel *k*, standard concentration results imply:

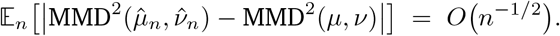

However, the dependence of this convergence rate on the dimension *d* crucially hinges on the choice of kernel and bandwidth. For the Gaussian kernel, an appropriately selected bandwidth—commonly chosen via heuristics like the median-distance heuristic or cross-validation—typically yields a dimension-independent convergence rate due to favorable exponential concentration properties ^43**?**^. Conversely, choosing an excessively small bandwidth (large *γ*) results in a kernel sharply peaked around zero, thus emphasizing pairwise distances and making the estimation more sensitive to dimensionality. In such cases, the empirical MMD estimation can deteriorate, potentially degrading toward a dimension-dependent rate 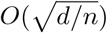.

### 2.3 Energy distance

An alternative choice for *k* is using a negative distance,

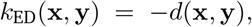

which is conditionally positive definite on ℝ^*d*^ and also characteristic. This ensures that the corresponding MMD can distinguish between distinct *µ* and *ν*. In practice, no bandwidth parameter is required, as the distance is fully determined by the Euclidean distance ∥**x** − **y**∥ itself.

By substituting *k*_ED_ into the MMD definition 13, we obtain

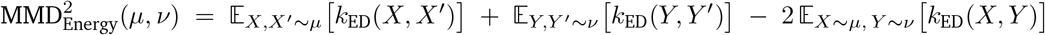

We follow Heumos et al. ^37^, Peidli et al. ^45^ in choosing *d* to be the Euclidean distance.

#### Sample Complexity

As discussed for MMD and *R*^2^, the empirical estimation of energy distance (*k*(*x, y*) = −∥ *x* − *y* ∥) initially converges at a parametric rate *O*(*n*^−1/2^) when the dimension *d* is fixed. However, unlike the Gaussian kernel used for MMD, energy distance does not have a bandwidth parameter to mitigate the effects of dimensionality. Thus, its convergence rate naturally degrades to 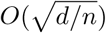 in higher dimensions, reflecting the difficulty of accurately estimating pairwise distances as *d* increases ^44^.

#### Advantages and disadvantages

Advantages of the energy distance are its fast computation and its independence of any hyperparameter. Moreover, it is a distributional metric, and hence takes into account all moments of the distribution.

Disadvantages are limited interpretability and the curse of dimensionality in high dimensions.

### 2.4 Sinkhorn divergence

As discussed in 1.3, OT provides a powerful method for comparing probability measures by minimizing a transport cost between them, but classical OT distances, such as the Wasserstein distance, can be computationally expensive and suffer from the curse of dimensionality. As explained in 1.3, a popular approach to mitigate these issues is to add an entropic regularization term, yielding the entropic Wasserstein distance. Concretely, the entropic Wasserstein distance is defined by

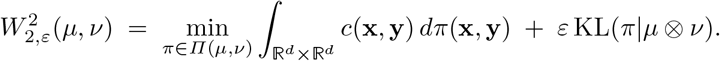

Although 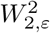 reduces to a valid distance when *ε* = 0, for *ε >* 0 it fails to be a valid divergence because 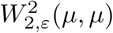 can be strictly positive. To address this, Feydy et al. ^46^, Genevay et al. ^47^ introduced the Sinkhorn divergence, which “de-biases” the entropic Wasserstein by subtracting self-interactions:

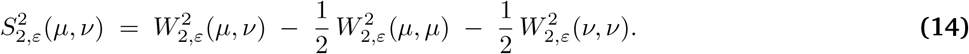

This correction ensures that 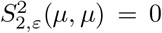 if and only if *µ* = *ν* ^46^ Theorem 1, thereby recovering the separation property that is lost in the entropic Wasserstein for *ε >* 0. Due to this property, and similarly to the MMD, the Sinkhorn divergence has found wide use in generative modeling ^7,29,47–49^.

#### Sample Complexity

In practice, and as for the MMD, we compute the Sinkhorn divergence on empirical distributions of cells 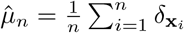 and 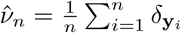, by evaluating

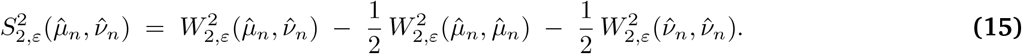

We compute each term using the Sinkhorn algorithm 1 (with *τ*_1_ = *τ*_2_ = 1, here). From a statistical perspective, the Sinkhorn divergence 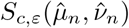 also enjoys a favorable sample complexity. More precisely,

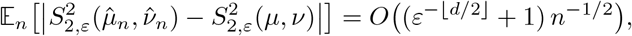

where 𝔼_n_ denotes expectation with respect to **x**_1_, …, **x**_n_ ~ _iid_ *µ* and **y**_1_, …, **y**_n_ ~ _iid_ *ν* ^20,23,50^. By tuning *ε*, this rate transitions smoothly from the classical *O*(*n*^−1/d^) complexity (as *ε* → 0, recovering unregularized OT) to the *O*(*n*^−1/2^) rate typical of MMD-like discrepancies (as *ε→ ∞*). In doing so, the Sinkhorn divergence sidesteps much of OT’s curse of dimensionality ^51,52^, while still retaining crucial geometric features of transport for moderate values of *ε*.

If not stated otherwise, we compute 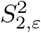 for *ϵ* ∈ 0.1, 1, 10 and report the mean.

#### Advantages and disadvantages

A drawback of the Sinkhorn divergence is its computational burden due to solving three entropic OT problems. Moreover, despite the possibility to set a hyperparameter in order to address the curse of dimensionality, it cannot avoid it completely. Additionally, it is not straightforward how to choose the entropic regularization *ε*.

Advantages of the Sinkhorn divergence include it being a distributional metric, thus taking into account all moments of the distribution. Additionally, it allows for a relatively straightforward intuition thanks to its motivation from an optimal transport perspective.

### 2.5 Evaluation of metrics

Having introduced the metrics, it remains to define which spaces the metrics are computed on.

If not stated otherwise, CellFlow generates its prediction in latent space (PCA or VAE), and generated cells are decoded to (log1p-transformed and normalized) gene space. This allows to compute metrics in gene space. To avoid low convergence rates due to the curse of dimensionality of distributional metrics, we only assess the R squared in gene space.

To be able to compute meaningful distributional metrics, we project predictions into a latent space, typically PCA-space between 10 and 100 dimensions, depending on the complexity of the dataset. This PCA space is normally different from the one which e.g. CellFlow was trained on, as PCA spaces constructed from CellFlow are computed from the training data only, while the space for evaluation is computed from the whole dataset to ensure that all axes of variation are captured. As a sidenote, we would like to point out that the Sinkhorn divergence in latent space is the most equivalent evaluation to the computer vision domain. In fact, generative models for images are typically evaluated by the Wasserstein distance between true and predicted images in the latent space of the Inceptionv3 model ^53^.

Finally, we follow recent literature ^38–41^ in computing the R squared in condition-specifically computed differentially expressed gene (DEG) space. In effect, for each condition, we compute the 50 top differentially expressed genes using scanpy.

## 3 Applications

### 3.1 PBMCs treated with cytokine

We downloaded the data from https://www.parsebiosciences.com/datasets/10-million-human-pbmcs-in-a-single-experiment/. It contains measurements of 90 cytokine treatments for PBMC samples from twelve donors, as well as control measurements for each of the donors. We filtered for 2000 highly variable genes using scanpy’s `highly_variable_genes` function, and performed normalization per cell (`normalize_total`) followed by log1pnormalization. Following the authors of the mouse cytokine dictionary ^54^, we computed differentially expressed genes (DEGs) using a Wilcoxon test specific to each cell type, donor and cytokine treatment.

For computing donor and cytokine similarities, we first computed the 2000-dimensional donor-specific mean response vector to each cytokine treatment. To obtain donor similarities, we computed cosine similarities between donor-specific response vectors computed as the concatenation (of size 90 · 2000) of mean response vectors. Analogously, for computing cytokine similarities, we computed cosine similarities between cytokine-specific vectors containing the concatenation (of size 12 · 2000) of mean response vectors. When considering cytokine family-specific donor similarities, we computed the cosine similarities between vectors containing the donor-specific concatenation of the mean responses of cytokines belonging to the cytokine family, i.e. concatenations of size *k ·* 2000, where *k <* 90 is the number of cytokines belonging to the cytokine family considered.

For the task of predicting the donor-specific effect of a (partially) unseen cytokine, we selected a random subset of cytokines (4-1BBL, ADSF, APRIL, BAFF, C5a, IFN-beta, IL-13, IL-15, Noggin, OSM, O**x**40L, IFN-epsilon) whose responses we predicted for all donors whose response to the test cytokine had not been seen during training in the respective split. For each of the 12 cytokines above, we created training sets with different sets of donors included (note that we always include some measurements for each donor, here, we refer to donors being included with respect to the twelve test cytokines), and predicted the donor-specific response for all remaining donors. In particular, given a test cytokine, we trained one model containing i=0 measurements of the cytokine (one model, twelve patient-specific distributions predicted). For each *i* = 1, …, 10, we created three sets with i donors included (for each i, three models trained, and for each model, 12 − *i* patient-specific distributions predicted). Moreover, for *i* = 11, we trained 12 models, with every single donor left out once. This resulted in 1 + 10 · 3 + 12 = 43 models for each cytokine, resulting in 12 · 43 = 516 trained models, and 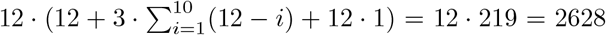 patient-cytokine-specific distributions.

Analogously, for assessing the predictive performance with respect to the number of included cytokines for donor-specific response modeling, we included different sets of cytokines in the training data for predicting the donor-specific effect of a fixed set of ten randomly chosen cytokines (FasL, IFN-omega, IL-1Ra, O**x**40L, CD27L, IL-32-beta, M-CSF, BAFF, IFN-gamma, ADSF). For each donor, we created training sets with different sets of cytokines (from the set of 90 − 10 = 80 cytokines whose responses are never predicted), and predicted the donor-specific response for all ten cytokines listed above. Given a test donor, we included three random subsets with *i* = {1, 2, 4, 8, 16, 32, 64} measurements of cytokines for this donor, while for all remaining donors we include all 90 measurements. Similarly, for each donor we trained one model with *i* = 0 cytokine treatments observed for that donor. On the other extreme, for each donor we trained one model with all 90 − 10 = 80 donor-specific cytokine responses in the training data set. This resulted in 12· (3· 7 + 1 + 1) = 12 · 23 = 276 trained models and thus 276 · 10 = 2, 760 donor-specific cytokine-treated populations which were predicted. We evaluated distributional metrics on a latent space computed by projecting predictions (living in log1p-transformed normalized count space) into 30-dimensional PCA space computed from the full dataset.

For computing cell type specific metrics, we transferred the labels using the label of the nearest neighbor (CellFlow’s `compute_wknn` and `transfer_labels` functions) of a reference dataset, which is a random subset of the full data (to accelerate the label transfer). We note that we assume a subset of the full data to be sufficient for cell type transfer due to the relatively small number of cell types in the dataset. All predictive models generate 10000 cells for a single distribution.

#### Identity

Given a donor and a cytokine, the predicted perturbed population is the control population of the given donor. Intuitively, this model is good for cytokines inducing a low effect, while it is bad for cytokines inducing a strong effect.

#### Mean test donor response across cytokines

Given a donor *d*_0_ with control population 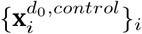, this mean model estimates the perturbed population to be the mean effect across all other cytokines for donor *d*_0_. Therefore, let 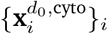 be the perturbed population for donor *d*_0_ and cytokine cyto ∈ 𝒞, with 𝒞 denoting the set of cytokines. Let 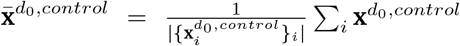 be the mean vector of the donor-specific control population. Then, the mean displacement vector for donor *d*_0_ and cytokine cyto_0_ is computed as 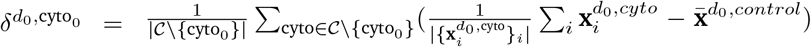. Thus, the population of cyto_0_-perturbed cells is modelled as 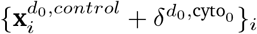.

#### Mean test cytokine response across donors

Given a donor *d*_0_ in the set of donors 𝒟 with control population 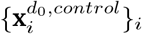, and a cytokine cyto_0_, this mean model estimates the perturbed population of donor *d*_0_ treated with cytokine cyto_0_ to be the mean effect of cyto_0_ across all other donors. Therefore, let 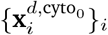 be the population of cells treated with cyto_0_ for donor 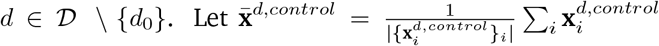 be the mean vector of the donor-specific control population of donor *d*. Then, the mean displacement vector of cyto_0_ is computed as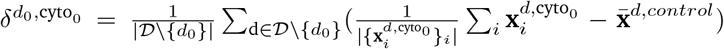. Thus, the donor-specific population of cyto_0_-perturbed cells is modelled as 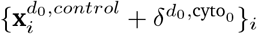.

#### CellFlow

The perturbation of CellFlow was set to be the cytokine treatment, which we embedded using ESM2 representations. There is no perturbation covariate. The sample covariate was the donor, which we one-hot encoded for the first task (Figure 1B), while for the second task (Figure 1F), we embedded donors using the mean of the (processed) gene expression of the control distribution. We trained CellFlow on 100-dimensional PCA space. For the first task, we trained the models in 100-dimensional cytokine/donor-specific PCA space. I.e., given a cytokine, the PCs were computed based on the full set of cytokine-treated cells (and control cells) which are not to be imputed. For the second task, given one test donor, we fitted the PCA-encoder using all cells measured for all remaining donors.

Due to the large number of considered tasks and conditions, we did not perform a comprehensive hyperparameter search. We thus conducted a non-systematic hyperparameter optimization on the task of predicting donor-specific IFN-beta treated populations with respect to the energy distance. Hyperparameter ranges as well as final model configurations deviating from the default configuration can be found in Supplementary Table 1.

We computed the scaling law using a linear fit in log2 space of the energy distance and log2 transformed number of cytokines seen for the test donor. R-squared values of the fitted curves across all numbers of cytokines are 0.916 for CellFlow and 0.618 for the mean model (mean cytokine response), while it is not defined for the other baseline models due to their constant performance. For the R-squared values fitted on the four data points corresponding to the four largest numbers of cytokines included in the training data set (16, 32, 64, 80), R-squared values are 0.999 for CellFlow and 0.980 for the mean model.

For computing feature importance scores for the CellFlow model, we fitted a LASSO model *y* ~ *β*_0_ + ∑_cyto_ *β*_cyto_ *I*_cyto_ + ∑ *k*∈ 𝒦 *β* _*k*_ *I* _*k*_ where *I*_c_ is the indicator function of cytokine cyto being included in the training data, and *I*_k_ being the indicator function of the CellFlow model containing *k* elements in the training dataset for the new donor. The target variable *y* is the energy distance. We optimize the hyperparameter of the linear model using sklearn’s `LassoCV` (using 5 folds), and report the coefficients *β*_cyto_ as feature importance scores.

### 3.2 Developing zebrafish

We obtained the dataset from the authors of ZSCAPE ^55^. We used the embedding `X_aligned` provided by the authors for all models, i.e. both CellFlow as well as the baseline models. For a detailed description of how the embedding was obtained, we refer the reader to the original manuscript. In short, a PCA-encoder (100-dimensional) was trained on the control cells, and perturbed cells were projected onto it, followed by integration in latent space. Importantly, the embedding was only generated from control cells, thus there is no data leakage from unseen perturbed cells. Due to the heterogeneity of the developing zebrafish, we were interested in distributional shifts rather than changes on single gene level, and thus all metrics were evaluated in the latent space `X_aligned`. Each predictive model generated 30000 cells.

#### Identity

Consistent with other use cases, the identity model assumes the perturbation to have no effect. As all other models, the identity model acts on the `X_aligned` space.

#### Mean across gene knockouts

Given a time point *t*_0_ with control population 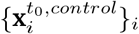, this mean model estimates the perturbed population to be the mean effect across all other genetic knockouts at the same developmental stage *t*_0_. Therefore, let 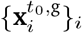 be the perturbed population at time *t*_0_ and gene knockout g, 𝒢 where 𝒢 is the set of gene knockouts. Let 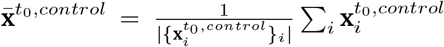 be the mean vector of the time point-specific control population. Then, the mean displacement vector for gene knockout *g*_0_ at time point *t*_0_ is computed as 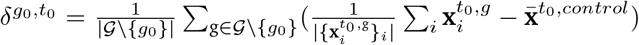. Thus, the population of perturbed vectors is computed as 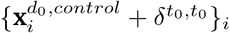.

#### Mean across time points

Given a gene knockout *g*_0_, this mean model estimates the perturbed population to be the mean effect of *g*_0_ across all other time points 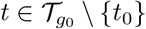 for which *g*_0_-mutants have been measured. Therefore, let 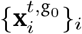 be the g_0_-perturbed population at time *t*. Let 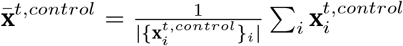 be the mean vector of the time point-specific control population. Then, the mean displacement vector for gene knockout *g*_0_ at time point *t*_0_ is computed as 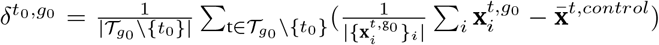. Thus, the population of *g*_0_-perturbed cells at time point *t*_0_ is computed as 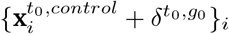.

#### CellFlow

We consider three different setups in this use case. The first one (Figure 2B-J) predicts a mutant’s phenotype at an unseen combination of time point *t* ∈ 𝒯 with 𝒯 = {18, 24, 36, 48, 72}[hpf] and genetic knockout g ∈ 𝒢.

The second one (Fig.2k) predicts a mutant’s phenotype at any 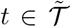 with 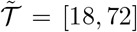 The third one predicts a mutant’s phenotype at all *t* ∈ 𝒯 for a completely unseen gene knockout g ∈ 𝒢.

We used the same hyperparameter configuration for all tasks. We did perform a random hyperparameter search with a budget of 300 runs on random held out conditions of gene knockout/developmental stage, see Supplementary Table 2 for parameters.

For the first task, we set up CellFlow with gene knockout(s) as perturbation with no perturbation covariate. Moreover, we encoded the log-transformed developmental stage as sample covariate. The source populations are control cells, i.e. given a time *t* and a genetic perturbation *g*, CellFlow learns a map from control populations at time *t* to the perturbed population under gene knockout *g* at time *t*. Parameters are provided in Supplementary Table 2. We trained 71 models, each one holding out one gene knockout/developmental stage combination. For computing true cell type proportions, we did not account for individual zebrafish, as we consider the task of learning the phenotype of a generic mutant rather than of individual mutants (for the latter there exists no ground truth). Instead, we computed cell type proportions from the union of set of cells corresponding to the combination of genetic perturbation and developmental time point, resulting in an ‘average mutant’. To quantify the variability of cell type proportions across individual zebrafish within the average mutant, we computed cell type proportion changes of individual zebrafish with respect to the average mutant. In particular, to have the same reference as the predicted perturbed population, the average mutant contains the cells of the individual mutant, resulting in a conservative estimate of the variability (Figure 3H,I).

For computing depletion scores of cdx4+cdx1a-perturbed phenotypes at time point 36, we built a joint graph of control and perturbed cells (once true, once predicted) using `rapids.pp.neighbors` with default parameters based on the `X_aligned` embedding. We then computed the fraction of neighbors which are perturbed cells, and test for the null hypothesis that the fraction of neighbors which are perturbed cells is not reduced. We conducted a permutation test using 1000 permutations of the neighborhood graph. The empirical p-value is the cell-specific depletion score.

For the second task (Figure 3K), equivalently to the first task, we set up CellFlow with gene knockout(s) as perturbation with no perturbation covariate. Moreover, we encoded the log-transformed developmental stage as sample covariate. However, the source distribution is only the control distribution at the earliest time point, i.e. at 18hpf. This allows to map from 18hpf to any other time point, for all perturbations, but also for the control condition (as e.g. 36hpf is a distribution which is mapped to). We focused on the development of the central nervous system (CNS), and thus subset the data to all cell types belonging to the CNS. We held out the time point 24hpf of cdx4+cdx1a-perturbed mutants to be able to compare the interpolated generated population. Subsequently, we generated populations corresponding to 18, 20, 22, …, 36hpf, transferred annotations using one-nearest neighbor, and finally computed cell type proportions analogously to task 1.

Finally, for the third task (Figure 3L,M), we setup CellFlow the same way as for the two previous tasks, but this time held out distributions for all 23 gene knockouts. We thus trained 23 models, each of which trained with a different mutant held out from the training dataset.

### 3.3 Sciplex data

We downloaded the sciplex-v3 data from pertpy (https://pertpy.readthedocs.io/en/latest/usage/data/pertpy.data.sciplex3_raw.html) and filtered for all conditions (i.e. combination of cell line, drug, and dosage) which contain at least 100 cells (for distributional metrics to be meaningful), resulting in 572,853 cells. As all models considered work on normalized counts, we used scanpy’s `normalize\_total` and `log1p` preprocessing functions before filtering for 2000 highly variable genes. We retrieved the SMILES using pertpy’s `annotate_compound` function, and did manual annotation of the drugs which were not found with `pertpy`. We obtained the Morgan fingerprints using `rdkit`’s ^9^ `MolFromSmiles`. We used the common benchmarking split introduced by chemCPA ^40^, i.e. the cells treated with Hesperadin, TAK-901, Dacinostat_(LAQ824), Givinostat_(ITF2357), Belinostat_(P**x**D101), Quisinostat_(JNJ-26481585)_2HCl, Alvespimycin_(17-DMAG)_HCl, Tanespimycin_(17-AAG), or Flavopiridol_HCl were not seen during training (i.e. all combinations containing any of these drugs are left out, regardless of the dosage or cell line).

#### Comparison of encoder-decoder methods

Importantly, CellFlow is encoder-decoder agnostic. Hence, an embedding has to be chosen. This should be done according to well-behavedness of the space for flow matching (e.g. a rather smooth than discretized space), while being performant in reconstruction. We compared PCA-embeddings with VAE embeddings for the latter. For the VAE embedding we adapted scVI’s JAX implementation ^56^ to a normal likelihood, as the input is log1p-transformed. We performed a hyperparameter search across the dimension of the latent space (10, 32, 64, 128, 256), and the number of hidden units per layer (512, 1024, 2048). We optimized the result with respect to the mean R-squared between decoded gene expression and true gene expression (log1p transformed) across the unseen perturbation conditions. The best configuration had 10 latent dimensions and 2048 hidden units per layer.

#### chemCPA

We had the authors of chemCPA optimize the hyperparameters of chemCPA in an updated implementation of the method, which we made available via our reproducibility repository. We note that no pretraining of chemCPA was performed.

#### CondOT

While the authors did not consider the use case of unseen drugs ^57^, but only predicted cellular responses to unseen dosages of seen drugs, we extended CondOT to work for this setting. In fact, we implemented PICNNs as described by the authors in Bunne et al. ^57^ using 4 hidden layers of width 64. For all networks, we use the Adam optimizer) with a learning rate of 0.0001 (*β*_1_ = 0.5, *β*_2_ = 0.9) and *λ* = 1. We performed a hyperparameter search across the following parameters: Number of PCs (50, 100, 300), number of pretraining iterations to learn the identity (see Eyring et al. ^58^, 0, 1,000, 10,000), number of iterations (50,000, 200,000), batch size (256, 1024).

#### biolord

The authors provide a hyperparameter configuration for exactly this use case, which we followed: https://github.com/nitzanlab/biolord_reproducibility/blob/main/notebooks/perturbations/sciplex3/2_perturbations_sciplex3_evaluation.ipynb.

#### CellFlow

Perturbations are encoded using the fingerprint emebdding described above, perturbation covariates are log10 dosages in nM. We embedded the cell line using embeddings obtained from the Cancer Cell Line encyclopedia ^11^, and input them as sample covariate. For representing cells, we used 100-dimensional PCA embedding. Parameters for hyperparameter optimisation (wrt to R squared in gene space, 300 runs) can be found in Supplementary Table 3.

### 3.4 Combosciplex data

We downloaded the data from pertpy (https://pertpy.readthedocs.io/en/latest/usage/data/pertpy.data.combosciplex.html). The data was preprocessed using scanpy’s `normalize_total` and `log1p` methods, followed by 2000 highly variable gene selection. As some of the competing methods can take count data as input, we then re-applied the normalisation procedure to the count data of the highly variable genes. Molecular fingerprints were calculated the same way as for the sciPlex dataset described above. We extracted all perturbed populations such that each single drug of the drug combination has been seen in at least one other perturbed condition, resulting in 27 (out of 31) perturbed populations, the remaining measurements, which are included in the training data of all splits, contain a drug which is not seen in any other measurement (Alvespimycin+Pirarubicin, Dacinostat+Danusertib, Givinostat+Carmofur, Givinostat+Tanespimycin). These 27 populations were randomly split into four splits, while ensuring that for each single drug appearing in a combination of drugs is also seen during training (in any combination).

CPA, chemCPA, and CellFlow are optimized with a random grid search with 300 trials with respect to the R squared in gene expression space. All models are optimized with respect to split 3.

#### CPA & chemCPA

We followed the tutorials for CPA (https://cpa-tools.readthedocs.io/en/latest/tutorials/combosciplex.html and chemCPA https://cpa-tools.readthedocs.io/en/latest/tutorials/combosciplex_Rdkit_embeddings.html). As these tutorials work on raw counts, while our evaluation is performed on normalized, log1p-transformed counts, we performed a hyperparameter search. In particular, we allowed the models to operate in both transformed and raw counts. In the latter case, the predictions were processed using scanpy’s `normalize_total` and `log1p` (the same processing which had been applied to).

#### Additive model

We computed displacement vectors for each treatment as the difference between the mean (normalized) gene expression of the perturbed population and the mean gene expression of the control population, i.e. for a perturbed population treated with drug *d*_0_, the displacement vector is computed as 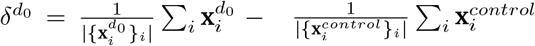 where 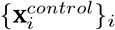 denotes the control population and 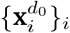 denotes the population treated with drug *d*_0_. As not all drugs which were applied in a combined treatment were also applied isolatedly, the additive model sometimes consists of multiple displacement terms. For example, given the combined treatment of drugs *d*_0_ and *d*_1_ *d*_0_ + *d*_1_, as well as the combined treatment *d*_1_ + *d*_2_, and the single treatment *d*_2_, we define the additive model for *d*_0_ to be 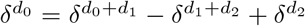, and thus the additive model for 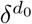 consists of three terms. Hence, the predicted population for drug *d*_0_ is 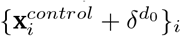. The definition of the additive model is provided in Supplementary Table 4.

#### CellFlow

Perturbations are drug treatments encoded as fingerprints. As all drugs are given with same dose, and all treatments are limited to one cell line, CellFlow doesn’t require any perturbation covariate or sample covariate. We used 100-dimensional PCA space as cell embedding (Supplementary Table 4).

Figure 4H-K are based on split 3. For fitting a regression model for predicting log fold-changes on the learn condition space of CellFlow (Figure 4J,K), we performed a hyperparameter search across different kernels (linear, polynomial, radial basis function, sigmoid) as implemented by `sklearn`, together with a range of hyperparameters (between 3 and 9 depending on the kernel), and performed a 3-fold cross validation with `GridSearchCV`. The linear kernel performed best, hence we reported its results.

### 3.5 Perturb-seq for gene overexpression

We downloaded the data from the link https://figshare.com/articles/dataset/perturbseq_nornan/22344253?file=39756463 which corresponds to the preprocessed dataset as provided by the authors of biolord ^41^. The dataset contains measurements of cells from the A549 cell line where either a single gene or a combination of two genes are perturbed. We follow the evaluation scheme as originally proposed by the authors in GEARS ^59^, and as subsequently applied by the authors of biolord. In line with this, the following perturbation conditions were excluded from the evaluation since GEARS is unable to make predictions for them as they are not part of its internal GO graph: ‘RHOXF2BB+ctrl’, ‘LYL1+IER5L’, ‘ctrl+IER5L’, ‘KIAA1804+ctrl’, ‘IER5L+ctrl’, ‘RHOXF2BB+ZBTB25’, ‘RHOXF2BB+SET’. The final preprocessed dataset contains 89357 cells (with 5045 genes) distributed across 101 single-gene and 128 double-gene perturbations. We use random data splits defined by the authors of biolord; in particular the dataset is split by perturbation condition into 5 random train-test partitions. Perturbation conditions in the test set can furthermore be divided into 4 categories: single-gene perturbation (single gene perturbation which was not part of the training data), double-gene-seen-2 (the combination of perturbed genes but both genes were included in the training data as single-gene perturbations), double-gene-seen-1 (only one of the genes was included as single perturbation in the training data) and double-gene-seen-0 (neither of the two genes was included as singlegene perturbation in the training data). The distribution of perturbation conditions across categories are: Split 1 contains 36 unseen single-gene perturbations, 12 unseen double-gene perturbations with none of the singletons seen, 44 double-gene perturbations with exactly one of the singletons seen, and 15 double-gene perturbations with both singletons seen. These numbers for split two are 37, 14, 51, 15; for split three 35, 13, 52, 9; for split four 37, 9, 53, 12; and for split five 40, 9, 55, 13.

We compute differentially expressed genes (DEG) between control and perturbed cells separately per condition and include the top 50 genes in the DEGs-based evaluation, analogously to the other tasks. Distributional metrics are evaluated in 10-dimensional PCA space. Analogously to all other tasks, this PCA embedding was computed on the full training dataset.

#### GEARS

We trained GEARS for 15 epochs using the optimal hyperparamenters reported in Supplementary Note 22 in the original publication ^59^. We used the `gears.inference.evaluate` to generate perturbed cells for the conditions in the test subset for each split.

#### scGPT

The scGPT evaluation closely follows the code (including all reported hyperparameters) of the authors which is available at https://github.com/bowang-lab/scGPT/blob/7301b51a72f5db321fccebb51bc4dd1380d99023/tutorials/Tutorial_Perturbation.ipynb. To make the code compatible with data saved with GEARS 0.0.4 and newer, we adapted the training function in the notebook and the prediction function (`pred_perturb`) in the scGPT codebase to infer the multi-hot perturbation encoding vector `pert_flags` from the perturbation index `pert_idx` instead of expecting it to come from the GEARS dataloader as done in https://github.com/const-ae/linear_perturbation_prediction-Paper/blob/15de7bf08a2f92a0511bafe0294b664b8e2e6ace/benchmark/src/run_scgpt.py#L232.

#### biolord

The biolord evaluation closely follows the code of the authors which is available at https://github.com/nitzanlab/biolord_reproducibility/blob/main/scripts/biolord/norman/base_experiment_norman.py using hyper-parameters that were optimized on this dataset (https://github.com/nitzanlab/biolord_reproducibility/blob/main/scripts/biolord/norman/norman_optimal_config.py). We note that instead of working with individual control cells, biolord uses the mean vector across control cells as input and predicts mean and variance per gene to characterize the perturbation response. We use the predicted means and variances to instantiate a Normal distribution per gene and sample 500 perturbation responses per condition. In contrast to only considering the mean response vector, this allows us to evaluate distribution-based metrics.

#### Identity model

We use the implementation of the identity model that was previously used by the authors of GEARS and biolord available at https://github.com/nitzanlab/biolord_reproducibility/blob/main/notebooks/perturbations/norman/1_perturbations_norman_preprocessing.ipynb.

#### Additive model

Following previous studies on this dataset ^60^, we define an additive model for the double-gene seen category. This is defined exactly the way it is computed for the combosciplex data, with the particular case that here, the displacement vector for the combined genetic perturbation *g*_0_ + *g*_1_ is always the sum of the displacement vectors of the indivual perturbation: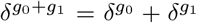, as the cell populations corresponding to the single genetic perturbations *g*_0_ and *g*_1_ have been observed.

#### CellFlow

Perturbations are genetic overexpressions encoded using ESM2 embeddings. There is no perturbation covariate, nor a sample covariate, as the experiment comprises only one cell line. We used 50-dimensional PCA representation as cell embeddings. We optimized the hyper-parameters of CellFlow on split 1.

### 3.6 Perturb-seq with pathway activations

We downloaded the data from https://zenodo.org/records/10520190 provided by the original publication ^61^. The datasets are provided as five Seurat subsets, one for each pathway, which we converted to `h5ad` format using the `sceasy` package and then merged using `anndata`. The data was filtered for perturbations with phenotypic effects using `mixscape` with a probability (perturbation score) threshold of 0.5 and at least 100 cells in both the source and target distributions. The filtered counts were normalized with scanpy’s `normalize_total` and `log1p` preprocessing functions before filtering for highly variable genes. Since the affected genes differed strongly between cell lines and cytokines, we calculated 500 highly variable genes per cell line/cytokines combination and used a joint set resulting in 8265 genes.

We considered three prediction tasks: Unseen genetic perturbations on seen cell line / cytokine (Figure S7A), seen genetic perturbations applied to an unseen combination of cell line and cytokine (but with the cell line and cytokine seen in different combinations, Figure S7C), and seen genetic perturbations to a completely new cell line (Figure S7F).

We performed a stratified 4-fold cross validation split of the genetic perturbations. In particular, we ranked the conditions based on their perturbation effect with respect to the R squared score for each cytokine separately (as the strength of perturbation can largely depend on the cytokine treatment). From each bin representing an effect size, we divided knockouts into four random groups and assigned them to the four different splits. This way, we obtained splits that are balanced in perturbation effect strength. The splits are available in the Supplementary Table 7.

For the second task, we selected five different combinations (including cell line BXPC3 with IFNG stimulation due to their close-up analysis in the original manuscript ^61^, and the remaining combinations randomly, but such that no cell line and no cytokine treatment is considered twice): BCPX3/IFNG, HAP1/TGFB, A549/IFNB, K562/INS, HT29/TNFA.

For the third task, we held out all conditions including the BXPC3 cell line.

We evaluated distributional metrics in 20-dimensional PCA space obtained from the full processed dataset. Perturbation effects for evaluation are with respect to R squared of condition-specific DEGs.

#### Identity model

Analogously to other tasks, the identity model predicts the perturbed population to be equal to the control population.

#### Mean model

The mean model is task-specific.

For task 1, we approximated the shift of a held-out knockout by the average shift from controls produced by the other (measured) knockouts, analogously to the mean models in other applications. Thus, the model assumes that the effect of a perturbation is constant for a combination of cell line and cytokine treatment.

For task 2 and 3, the genetic perturbations are modeled to be constant across cell lines, i.e. the mean displacement vector is computed from the same genetic knockout observed in different cell lines (as the gene knockouts are specific to cytokines, the measurements on cell lines induced with different cytokines are not taken into account).

#### CellFlow

We set up CellFlow with genetic knockouts as perturbations (without perturbation covariate) and cytokine treatment as sample covariate (due to the absence of control state, i.e. all cell lines have been treated with a cytokine). We embedded gene knockouts and cytokines with ESM2. For cell line embeddings, we obtained bulk RNA-seq data from the Cancer Cell Line Encyclopedia for each cell line and performed PCA with 100 principal components. Details are provided in Supplementary Table 6.

For the downstream analysis for task 2 (Figure S7E), we used the perturbation programs provided by the authors in Supplementary Table 3 of the original publication ^61^ and chose two programs that were especially highlighted in the original publication. For the left-out conditions, we generated the same number of cells per condition as in the ground truth distribution. Then, z-scores between the predictions and cell lines were calculated as follows:

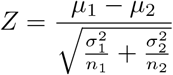

For the mean baseline model in Figure S7E, we calculated the mean of all gene-specific z-scores of the genetic perturbations on other cell lines. To quantify the similarity of the models (prediction or mean baseline model) and ground truth effects of each cell line, we computed the Pearson correlation of the z-scores of the target genes.

### 3.7 4i data

It is important to note that we learn one model per drug and the task consists of predicting the effect to unseen cells in contrast to predicting the effect to unseen treatments, as in all other use cases of this manuscript. We downloaded the 4i data from (ETH drive). We randomly selected 60% of the cells to be in the training set and 40% in the test set. Following the CellOT manuscript ^42^ we selected the same 48 of the in total 78 4i features, and also only used the same set of drugs the same drugs as the authors of CellOT.

#### scGen

We ran scGen ^38^ with default parameters following the procedure outline here.

#### CellOT

We ran CellOT with default parameters as given in the repository, with the only change being to replace the best model selection with respect to the test data by the best model selection with respect to the train data to prevent data leakage.

#### CellFlow

We used a constant dummy token as condition, and reduced the size of the architecture due to the smaller number of cells and smaller number of conditions. As we only observe one condition per model, we do not need to perform gradient averaging. This and other modifications to the architecture are described in Suppplementary Table 7.

### 3.8 NGN2-induced neuron morphogen screen

The dataset, including the annotations of cell type and regional identity, was provided to us by the authors of the original publication ^62^. Unannotated cells were filtered out and 4000 highly variable genes were selected using Seurat v3 algorithm ^63^. The total number of counts per cell was normalized to 10000, and the data was log-transformed. All the data processing was performed in scanpy ^36^.

#### Holdout splits

To evaluate performance on prediction combinatorial treatment outcomes, we constructed five holdout splits by withholding all experimental conditions containing certain combinations of AP and DV modulators in each split. Specifically, training sets always included cells from all single-molecule treatments and all conditions produced by half of all the molecule combinations. To ensure that performance differences across treatment combinations were not influenced by the over- or underrepresentation of individual treatments, we maintained a consistent proportion of every single treatment across all the random train-test splits. For example, among the four two-molecule combinations excluded, two always included BMP4, two included SHH, and each of the molecules acting on the anterior-posterior axis appeared once.

#### CellFlow training

For training the model, we used 30-dimensional PCA space. The embedding used for training was always constructed using only cells from the training set. All of the morphogen signaling modulators were one-hot encoded and the one-hot encodings multiplied by the corresponding dosages were used as input perturbation representations.

#### Baselines

As a baseline for any combination of treatment outcomes, we defined two baseline models:

- **Union**. Here, we computed the union of cells from the individual treatment conditions. This baseline assumes that a combination of treatments will result in the combination of states produced from individual treatments.
- **Barycenter**. Here, we computed the Wasserstein barycenter ^64^ between the distributions produced by individual treatments using the `FreeWassersteinBarycenter` implementation in `ott-jax` with a `Sinkhorn` solver and a fixed *ϵ* of 0.1. This baseline assumes that a combination of treatments will result in intermediate states between states produced from individual treatments.

#### Hyperparameter optimization

We performed hyperparameter tuning over a fixed set of hyperparameter values using the Bayesian optimization-based method Optuna ^65^. For CellFlow, a sweep was optimized to minimize E-distance on a held-out test set comprising the combinatorial treatments CHIR+BMP4 and FGF8+SHH. These conditions were excluded from any subsequent model evaluations. The model with the best MMD was used for evaluation. Biolord and CPA were hyperparameter-tuned to minimize MMD on the same holdout split using a grid of hyperparameters of comparable size to that of CellFlow. Each run of Optuna included between 400 and 600 of trained models. All optimized hyperparameters are reported in Supplementary Table 8.

#### Evaluation

For all of the models, a full gene expression matrix was reconstructed from the obtained predictions, with subsequent projection to the 20-dimensional PCA space of the whole ground truth dataset for evaluation. In addition to computing distributional metrics, predicted cells were assigned to the ground truth data clusters using weighted k-nearest neighbors, as described for the annotation of the human neural organoid cell atlas based on the primary atlas ^66^. We then computed the cosine similarity between predicted (*x*) and ground truth (*y*) cluster compositions as

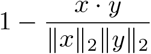

To visualize predictions, we projected them onto the UMAP embedding computed from the full dataset using `UMAP.fit` and `UMAP.transform` from cuML (https://github.com/rapidsai/cuml).

#### Morphogen interaction score

To analyze interactions between treatmetns and to assess whether combinations result in new cell states, we computed the MMD between the cells from combinatorial condition, either predicted or ground truth, and the union of cells from the corresponding single-treatment conditions.

### 3.9 Brain organoid morphogen screens

To train and evaluate CellFlow for predicting organoid engineering outcomes, we obtained three scRNA-seq datasets of organoid morphogen screens. The Amin dataset ^67^ was obtained from the link provided in the publication, while the other two datasets ^68,69^ (Azbukina and Sanchis-Calleja) were provided by the authors. All datasets were consistenly preprocessed using scanpy ^36^.

**Integration and annotation** To integrate all three datasets, we used HNOCA-tools (https://github.com/devsystemslab/HNOCA-tools) and a scANVI model ^70^ that was previously pre-trained on an atlas of the developing human brain ^66,71^. We performed query-to-reference mapping using the model on the organoid datasets using the datasets label as the batch covariate. We trained the model with

- `retrain = “partial”`
- `max_epochs = 200`
- `batch_size = 1024`
- `weight_decay = 0.0`

and otherwise default parameters. This shared latent space with the three organoid datasets as well as the primary cell atlas also allowed us to obtain consistent brain region and cell class annotations for all organoid datasets. For this, we used CellFlow’s `compute_wknn` and `transfer_labels` functions to transfer labels as described previously ^66^.

#### Holdout splits

To evaluate different aspects of model performance, we constructed two types of holdout splits:

- **Combination prediction**. Here, we sought to test the ability of the model to predict the outcome of unseen combinatorial morphogen treatments. For this, we selected morphogen combinations where conditions were available for both individual and combinatorial treatments, resulting in 12 separate splits. During each training run, we included individual treatments but held out their combination and all conditions including this combination from training.
- **Information transfer**. Here, we tested to what degree a condition seen in one dataset could inform predictions on other datasets. For this, we selected morphogens that were used in all three datasets, but in no more than 30% of conditions for each dataset, resulting in 7 separate splits. During each training run, we held out all conditions including this morphogen in one dataset from training.

All splits for both tasks are listed in Supplementary Table 9. For each holdout split, we retrained the scANVI model on the training data only, to obtain the latent embedding for training CellFlow and for computing baseline models.

#### CellFlow training and digital protocol encodings

We trained CellFlow on all thee combined datasets using the shared latent space of the scANVI model, which was recmputed for each split. To construct conditions for training, we obtained digital protocol encodings as follows: For each morphogen treatment, the morphogen was one-hot encoded and the one-hot encoding multiplied by the corresponding log-transformed dosages. We concatenated this representation of the morphogen with the start and end time of treatment (in days) as well as a one-hot encoding of the modulated pathway and the directions of modulation (−1 for inhibition, 1 for activation). Each condition was represented as a combinatorial set of such treatments and the dataset label was used as a sample covariate. Because not all datasets had controls that were not treated with any morphogen, we derived a common source distribution of 10000 cells by repeatedly computing the mean of 10 cells sampled from the primary atlas of the developing human brain. We trained CellFlow to generate predictions from this source distribution for all datasets based on the set of treatments and the dataset covariate.

#### Baselines

As a baseline for combinatorial treatment prediction we used the Union and Barycenter baselines as described above. For the information transfer task, we used an additional baseline:

- **Train dataset**. Here, we used all conditions in the respective dataset excluding the held-out test conditions as a baseline. This baseline assesses whether the model learns relevant treatment-specific effects beyond the effects of the base protocol used in each dataset, i.e. the dataset covariate label.

#### Hyperparameter optimization

We performed hyperparameter tuning over a fixed set of hyperparameter values using the Bayesian optimization-based method Optuna ^65^. We optimized for 200 iterations to minimize MMD on a holdout split where all conditions including BMP4 in the Sanchis-Calleja dataset ^69^ were held out during training. All optimized hyperparameters are reported in Supplementary Table 10.

#### Evaluation

For evaluation of CellFlow and baseline models, we computed distributional metrics directly in scANVI latent space as described above. In addition to computing distributional metrics, predicted cells were assigned to the ground truth data clusters using WKNN-based label transfer from the full dataset in scANVI latent space and cosine similarity was computed on these cluster compositions as described above. To visualize predictions, we projected them onto the UMAP embedding computed from the full dataset using `UMAP.fit` and `UMAP.transform` from cuML (https://github.com/rapidsai/cuml).

#### Virtual neural organoid protocol screen

To perform a virtual organoid protocol screen, we first composed protocols based on single, double and triple combinations of 14 morphogen pathway modulators as well as their concentration and timing. For all morphogens we fixed the concentration to the highest concentration observed across all datasets. To comprehensively probe the space of possible protocols without ending up with intractably many configurations, digital protocols were computed in two parts:

1. First, we considered four timings (early [1, 8], mid [8, 15], late [15, 22] and early-late [1, 22]) and 9 morphogen pathway modulators (SHH+PM, SAG, FGF2, FGF8, BMP4, BMP7, LDN, CHIR, RA). From these we computed all possible combinations of 1-3 morphogens each applied in all possible time windows.
2. Next, we considered six timings (early [1, 8], mid [8, 15], late [15, 22], early-mid [1, 15], mid-late [8, 22] and early-late [1, 22]) and 14 morphogen pathway modulators (SHH+PM, SAG, FGF2, FGF8, FGF19, FGF17, BMP4, BMP7, LDN, CHIR, RA, CycA, XAV, Insulin). Here, we computed all possible combinations of 1-3 morphogens but constrained timings such that all morphogens in a combination were given during the same time window.

After combining both parts, we ended up with 7951 unique protocol configurations. We further combined each of these protocols with one of the three dataset labels (i.e., the “base protocol”), which resulted in 23853 total digital protocols. We used CellFlow to generate predictions for each of these protocols based on 3000 cells sampled from the source distribution. This resulted in the virtual organoid protocol screen dataset comprising 70’315’442 cells. We annotated this dataset with brain region, cell class and cluster labels using WKNN-based label transfer from the human developing brain atlas.

To analyze the association between morphogen treatment and brain region composition across base protocols in the virtual screen, we focused on all protocols with single and double combinations of morphogens. We for each brain region, fitted a LASSO-regularized linear model as

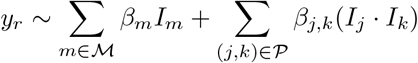

where *y*_r_ is the proportion of cells belonging to brain region *r*, ℳ is the set of all morphogens, *I*_*m*_ is the indicator variable for the presence of morphogen *m* and 𝒫 is the set of all pairwise combinations of morphogens. We used the `cv.glmnet` function from the glmnet R package ^72^ where the optimal regularization parameter *λ* is selected through cross-validation. All non-zero coefficients were treated as significant associations between morphogen treatment and region proportion.

To characterize the generated protocols, we computed two complementary metrics for each protocol:

- To assess how realistic the generated cell distributions were, assessed the similarity to cell states in the human developing brain atlas as

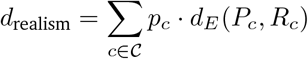

where *d*_*E*_ represents the energy distance, 𝒞 is the set of all reference clusters, *p*_c_ is the proportion of cells belonging to cluster *c, P*_c_ is the predicted distribution for cell cluster *c, R*_c_ is the corresponding distribution of the cluster in the reference atlas.
- To assess the novelty of generated cell distributions, we assessed the similarity to conditions in the training dataset as

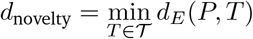

where *d*_E_ represents the energy distance, *P* is the predicted cell distribution, 𝒯 is the set of all conditions in the training data and *T* is the distribution corresponding to a single training condition.

